# Distinguishing shadows from surface boundaries using local achromatic cues

**DOI:** 10.1101/2022.03.08.483480

**Authors:** Christopher DiMattina, Josiah Burnham, Betul Guner, Haley Yerxa

## Abstract

In order to accurately parse the visual scene into distinct surfaces, it is essential to determine whether a local luminance edge is caused by a boundary between two surfaces or a shadow cast across a single surface. Previous studies have demonstrated that local chromatic cues may help to distinguish edges caused by shadows from those caused by surface boundaries, but the information potentially available in local *achromatic* cues like contrast, texture, and penumbral blur remains poorly understood. In this study, we develop and analyze a large database of hand-labeled achromatic shadow edges to better understand what image properties distinguish them from occlusion edges. We find that both the highest contrast as well as the lowest contrast edges are more likely to be occlusions than shadows, extending previous observations based on a more limited image set. We also find that contrast cues alone can reliably distinguish the two edge categories with nearly 70% accuracy at 40×40 resolution. Logistic regression on a Gabor Filter bank (**GFB**) modeling a population of V1 simple cells separates the categories with nearly 80% accuracy, and furthermore exhibits tuning to penumbral blur. A Filter-Rectify Filter (**FRF**) style neural network extending the **GFB** model performed at better than 80% accuracy, and exhibited greater sensitivity to texture differences. Comparing the models with humans performing the same occlusion/shadow classification task using the same stimuli reveals better agreement on an image-by-image basis between human performance and the **FRF** model than the **GFB** model. Taken as a whole, the present results suggest that local achromatic cues like contrast, penumbral blur, and texture play an important role in distinguishing edges caused by shadows from those caused by surface boundaries.

**AUTHOR SUMMARY:** Distinguishing edges caused by changes in illumination from edges caused by surface boundaries is an essential computation for accurately parsing the visual scene. Previous psychophysical investigations examining the utility of various locally available cues to classify edges as shadows or surface boundaries have primarily focused on color, as surface boundaries often give rise to a change in color whereas shadows will not. However, even in grayscale images we can readily distinguish shadows from surface boundaries, suggesting an important role for achromatic cues in addition to color. We demonstrate using statistical analysis of natural shadow and occlusion edges that locally available achromatic cues can be exploited by machine classifiers to reliably distinguish these two edge categories. These classifiers exhibit sensitivity to blur and local texture differences, and exhibit reasonably good agreement with humans classifying edges as shadows or occlusion boundaries. As trichromatic vision is relatively rare in the animal kingdom, our work suggests how organisms lacking rich color vision can still exploit other cues to avoid mistaking illumination changes for surface changes.

## INTRODUCTION

Luminance edge detection is an essential visual computation for identifying the boundary between two surfaces (**Marr, 1982**; **Martin, Fowlkes & Malik, 2004; DiMattina, Fox, & Lewicki, 2012; Mely, Kim, McGill, Gou, & Serre, 2016**). However, it is well known that in natural images, luminance edges may arise from several causes other than surface boundaries: Specular reflections, changes in material properties, changes in surface orientation, and cast shadows all can give rise to changes in luminance that do not correspond to surface boundaries (**Ramachandran 1988; Mamassian, Knill, & Kersten, 1998; Vilankar, Golden, Chandler, & Field, 2014; Casati & Cavanagh, 2019**). Shadows are one class of luminance edge which are essential to distinguish from other potential causes, since shadows do not indicate a change in surface reflectance properties, but simply a change in illumination (**Olmos & Kingdom, 2004**; **Kingdom, 2008, 2019**).

Recent work has addressed the question of how the visual system distinguishes material or reflectance edges from shadow edges in natural vision (**Breuil, Jennings, Bartheleme, Guyader, & Kingdom, 2019**). In their study, human observers classified both color and grayscale image patches viewed through an aperture as either shadow edges or material edges. These authors found that observers performed significantly better with the color images, and that a machine classifier making use of chromatic cues outperformed a machine classifier only using luminance cues. These findings are consistent with several studies suggesting an important role for color in distinguishing illumination from material (reviewed in **Kingdom, 2008, 2019**). In addition to improved performance with chromatic information, these investigators also observed improved performance with increasing aperture size. This may be because at larger spatial scales, texture information becomes available which is not available at smaller spatial scales.

Here we address the question of what locally available *achromatic* information is available to distinguish surface occlusion boundaries from cast shadows. This question is important since distinguishing material properties from the illuminant is a universal problem that all visual systems face, yet trichromacy is relatively rare in the animal kingdom. The vast majority of mammals are dichromats, and some mammals (for instance, owl monkeys) are actually cone monochromats. For such organisms, chromatic cues will be less useful, or even completely unavailable, for shadow segmentation. Furthermore, even in trichromatic species, color information is by no means necessary to segment shadows, as a casual inspection of the grayscale images in **Fig. 1a** reveals. This question is also important for the development of computer vision methods for shadow segmentation and removal in grayscale images. Most algorithms developed for this purpose utilize color information to distinguish changes in illumination from changes in surface reflectance (e.g., **Olmos & Kingdom, 2004**). However, in some computer vision applications chromatic information may be unavailable, for instance when processing archival images which pre-date color photography.

**Figure 1:**
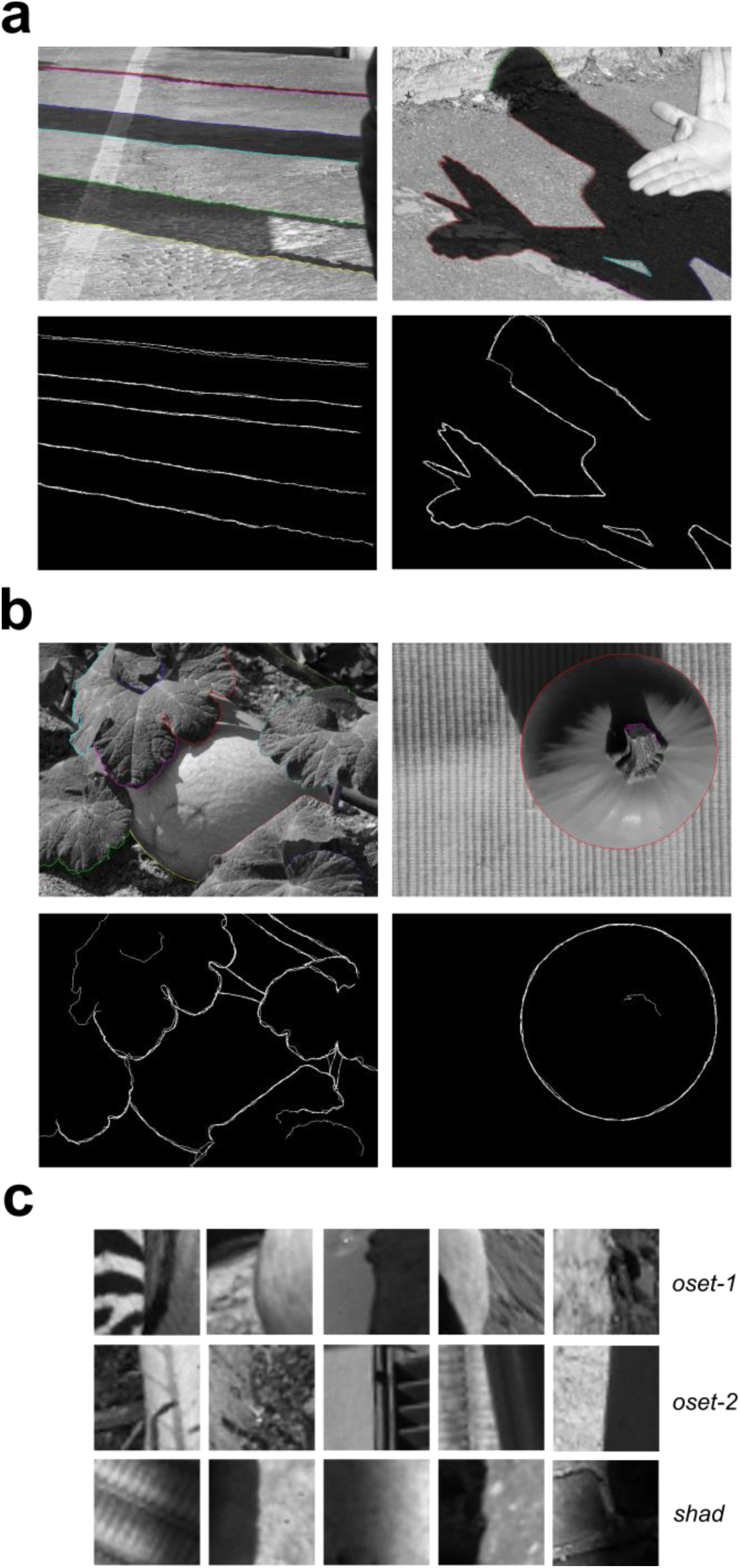
Images from the shadow boundary database and one occlusion database. (a) *Top*: Two representative images from our shadow database (**shad**) with shadow edges labeled by one observer. *Bottom:* Overlaid shadow boundaries obtained from all N = 4 observers. (b) *Top:* Two representative images from **oset-2**, with occlusions labeled by one observer. *Bottom:* Overlaid occlusion boundaries from all N = 4 observers. (c) Representative 40×40 image patches from each image set.

At least three possible sources of information are potentially available: (1) Differences in luminance contrast, (2) Differences in spatial luminance profiles, and (3) Texture cues. Pursuant to (1), quantitative Bayesian classifier analyses demonstrate that on average occlusions can be distinguished from non-occlusions (with shadows being one kind of non-occlusion edge) using only Michaelson contrast (**Vilankar et al., 2014**). However, in this study a fairly limited set of images was used, and it did not focus specifically on shadows. Pursuant to (2), it is well known that the luminance edges arising from shadows are often more gradual than those arising from surface boundaries due to penumbral blur (**Casati & Cavanagh, 2019; Vilankar et al., 2014**), and furthermore human observers can reliably distinguish sharp and blurred edges (**Watson & Ahumada, 2011**). Pursuant to (3), although texture segmentation is extremely well studied both psychophysically and computationally (**Landy & Graham, 2004; Victor, Chubb & Conte, 2017**), to our knowledge no psychophysical work has directly considered the role of local texture cues for classifying shadow and occlusion edges.

To address these questions, we develop a publicly available database of shadow edges by hand-labeling a set of N = 47 natural images from the McGill Color Calibrated Image Database (**Olmos & Kingdom, 2004**). We compare basic contrast and spatial frequency statistics of these shadow edges to the same properties measured from a novel occlusion database derived from a subset of 20 of the 47 images, as well as a previously published occlusion database containing 100 images (**DiMattina et al., 2012**). We demonstrate that machine classifiers defined using only 2 or 3 statistics can reliably discriminate 40×40 patches containing shadows from patches containing surface boundaries at nearly 70% accuracy when generalizing to images not used for training. We then evaluated the ability of an image-computable model comprised of a population of multi-scale Gabor filters resembling V1 simple cells to distinguish these two categories, attaining nearly 80% accuracy when generalizing to images not used for training. Furthermore, we found that our Gabor filter bank (**GFB**) model exhibited clear tuning to penumbral blur, a cue which often accompanies shadows. To test the hypothesis that local texture cues may be used to distinguish shadows from occlusion boundaries, we next trained a Filter-Rectify-Filter (**FRF**) model (**Landy & Graham, 2004; Zavitz & Baker, 2013, 2014; DiMattina & Baker 2019, 2021; DiMattina, 2022**) to solve the classification problem. Our best **FRF** model could classify 40×40 image patches taken from novel images at better than 80% accuracy on average. Analysis of the hidden units in these trained models demonstrated that many represented spatial differences in texture. Additional testing of the **FRF** model revealed reduced data-fitting performance with the removal of texture information, further underscoring the importance of local textures cues.

To compare human performance with that of machine classifiers, we performed a simple psychophysical experiment where observers classified 40×40 grayscale image patches as occlusion or shadow edges. We found that the best human observers performed about as well as the machine classifiers (**GFB**, **FRF**) performed generalizing to novel images (80% accuracy). By correlating human and model classification performance on an image-by-image basis, we observed a stronger correlation between human behavior and the **FRF** model predictions than the **GFB** model, even though the performance of the **FRF** model (as measured by percent correct) was only slightly better than the **GFB** model. As the **FRF** model exhibits a stronger sensitivity to texture cues, these results suggest that human observers are making use of textural cues to classify luminance changes as shadows or occlusions.

Overall, our results show that there are multiple, locally available achromatic cues which the visual system can potentially exploit to distinguish luminance changes arising from shadows from those caused by occlusion boundaries. It is of great interest for future research to explore the extent to which human observers utilize these various cues and how they combine with each other, and we suggest future psychophysical experiments motivated by our current results.

## RESULTS

### Shadow edge database

Four observers hand-labeled all of the shadow boundaries in a set of N = 47 images (786 x 576 pixel resolution) taken from the McGill calibrated color image database (**Olmos & Kingdom, 2004**). Two images from our shadow database (**shad**), together with the shadow edges annotated by one observer, are shown in the top panels of **Fig. 1a.** A binary overlay of the labels obtained from all four observers is shown in the bottom panels of **Fig. 1a.** To compare the statistical properties of shadow edges with occlusion edges, we utilized a previously developed database (**DiMattina et al., 2012**) of hand-labeled occlusion boundaries taken N = 100 images from the McGill Dataset (**oset-1**). In addition to **oset-1**, we also developed a second occlusion database (**oset-2**) from a subset of shadow database images (20 out of 47 images) by having the same four observers hand label occlusions in these images.

Two images from **oset-2** along with their hand-labeled occlusions are shown in **Fig. 1b**, and examples of image patches randomly selected from each dataset can be seen in **Fig. 1c**. The total number of hand-labeled occlusions and shadows in these images is quite large, with the **shad** database containing a total of 272,043 labeled shadow patches in 47 images. **oset-1** contains over 1.25 million occlusion patches (1,258,000), and **oset-2** contains 84,112 occlusion patches. All three databases (**shad**, **oset-1**, **oset-2**) are freely available to the scientific community at: https://www.fgcu.edu/faculty/cdimattina/.

### Analysis of image patch statistics

#### Contrast statistics

Previous investigators have characterized some of the statistical properties of occlusion edges and non-occlusion edges (including shadows) in natural scenes (**Vilankar et al., 2014**; **Breuil et al., 2019**). To connect with these previous explorations, we measured some of these quantities from both the occlusion and shadow edges: (1) RMS contrast 𝑐_𝑅𝑀𝑆_ (**Eq. 2**), and (2) Michelson contrast 𝑐_𝑀_ (**Eq. 3**). All analyses were performed on 40 x 40 pixel (5% of larger image dimension) image patches from one of our databases (**shad**, **oset-1**, **oset-2**), with the constraint that all patches were centered on a human-labeled edge (shadow or occlusion).

**Fig. 2a** plots the overlaid probability distributions of these quantities for the shadow images (magenta curves) and both occlusion databases (**oset-1**: *top*, **oset-2**: *bottom*, green curves). We see that the distribution of Michelson contrast (𝑐_𝑀_) for occlusions (green lines) spans the entire range of observed values, with occlusion edges giving rise to both the highest and lowest values. Furthermore, we see that occlusion edges are much more likely than shadow edges (magenta lines) to have very low values of 𝑐_𝑀_. We obtained median values of 𝑐_𝑀_ = 0.51 for our shadow set, and 𝑐_𝑀_= 0.30, 0.33 for **oset-1** and **oset-2**, respectively. Median values are significantly different between shadows and both occlusion sets (rank-sum test, *p* < 0.001). Similar results were obtained for RMS contrast (𝑐_𝑅𝑀𝑆_), although in this case the discrepancy between categories was smaller, and the distribution for occlusions was closer in shape to a normal distribution. For RMS contrast, we obtained median values of 𝑐_𝑅𝑀𝑆_= 0.64 for the shadows and 𝑐_𝑅𝑀𝑆_= 0.47, 0.59 respectively. Median values are significantly different between shadows and both occlusion sets (rank-sum test, *p* < 0.001). Large and highly significant positive correlations (Pearson’s *r*) were observed between these two contrast measures for all datasets (**oset-1**: *r* = 0.754, *p* < 0.001; **oset-2**: *r* = 0.722, *p* < 0.001, **shad**: *r* = 0.743, *p* < 0.001).

**Figure 2:**
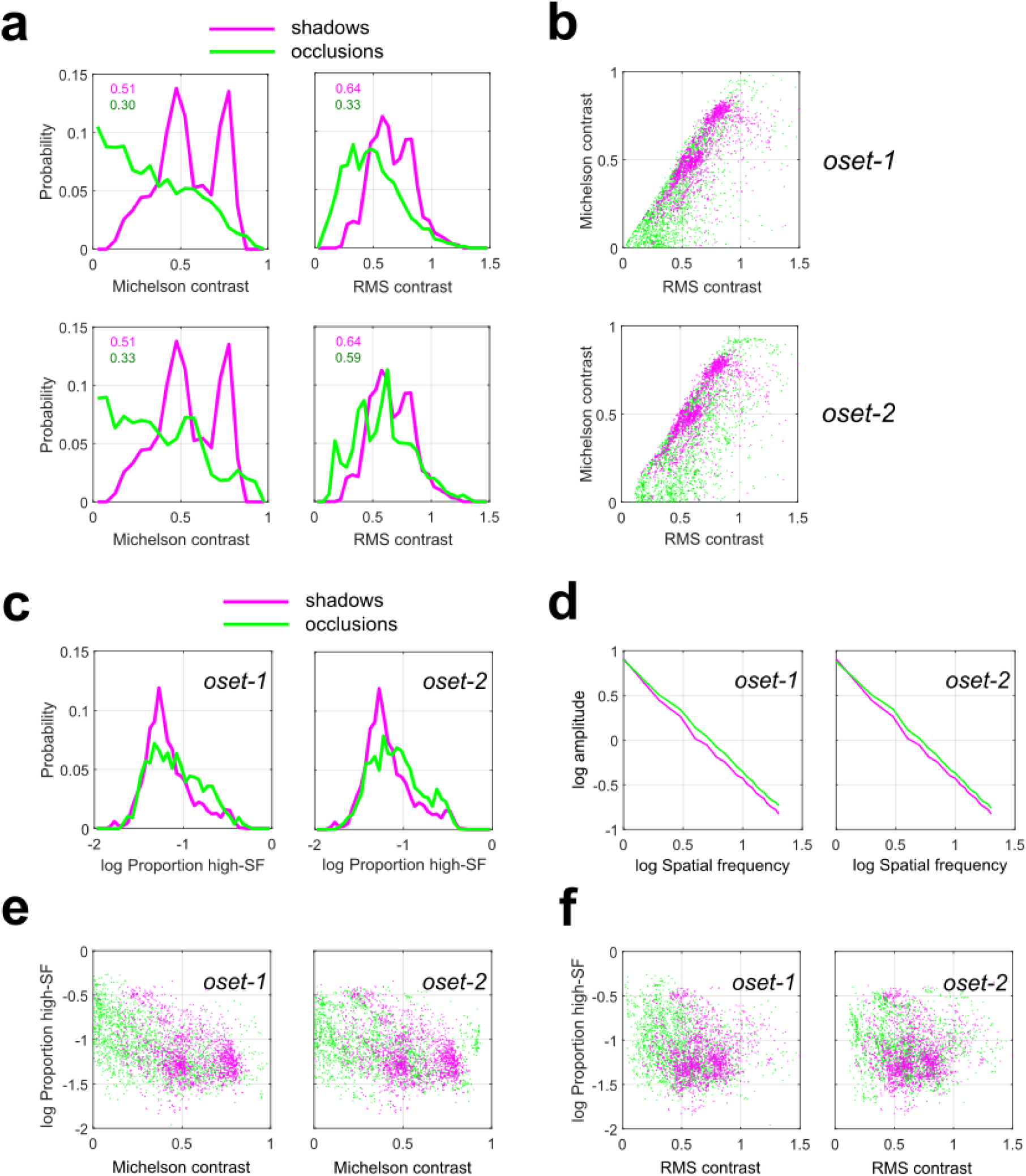
Contrast statistics measured from shadow and occlusion edges (N = 2000 of each category). Green curves/symbols indicate occlusions, Magenta curves/symbols indicate shadows. (a) Distributions of Michelson contrast (*left*) and RMS contrast (*right*) for our shadow database and two sets of occlusions (**oset-1**: *top*, **oset-2**: *bottom*). Note that for both occlusion sets, the highest and lowest contrast edges are occlusions. (b) Two-dimensional scatterplot of contrast measurements for shadow and occlusion edges for both sets of occlusions. The dashed line indicates the optimal separating hyperplane learned by a logistic regression classifier trained on a different set of observations. (c) Probability distribution of stimulus power 𝜋_ℎ_ in the high spatial frequency (>10 cycles/image) range for our shadow database (magenta curves) and both occlusion sets (green curves). N = 2000 for each set of images. (d) Rotational average of the amplitude spectrum for shadows (magenta curves) and both occlusion sets (green curves). (e) Scatterplots of 𝜋_ℎ_and Michelson contrast 𝑐_𝑀_for shadows (magenta symbols) and both occlusion sets (green symbols). (f) Same as (c) but for RMS contrast 𝑐_𝑅𝑀𝑆_.

Our finding that the lowest-contrast edges (by both measures) were occlusions rather than shadows is consistent with the fact that occlusion edges can often arise from two surfaces having identical mean luminance (**DiMattina et al., 2012**), whereas by definition a shadow edge requires a change in luminance. However, in agreement with previous measurements using a different set of occlusion patches identified using different methodology (**Vilankar et al., 2014**), we see that the far-right tails of the contrast distributions are dominated by occlusions, meaning that the highest contrast edges are more likely to be occlusions rather than shadows. Since 19 of the 38 images analyzed by **Vilankar et al. (2014)** were also contained in **oset-1**, we repeated our analysis on this subset of images, finding results consistent with our analyses based on the full image set (**Supplementary Fig. S1**).

We also see from **Fig. 2a** that in our sample the distribution of 𝑐_𝑀_ (but not 𝑐_𝑅𝑀𝑆_) appears to have some degree of bimodality. Most likely this reflects the properties of our database rather than the physics of shadow formation, as our shadow set contained a sub-set of images taken in conditions of strong illumination from a point source (the sun). Comparing the green curves on the top and bottom of **Fig. 2a**, we see a strong qualitative consistency between both sets of occlusion images. This provides us with reassurance that our measurements of contrast statistics reflect the general properties of occlusion edges rather than the idiosyncrasies of our databases.

**Fig. 2b** shows scatterplots of our measured quantities (𝑐_𝑀_, 𝑐_𝑅𝑀𝑆_) for both occlusion (green symbols) and shadow edges (magenta) for both sets of occlusions. For **oset-1**, a classifier using these two cues can separate occlusions and shadows with an average performance of 67.7% accuracy on test image sets not used for classifier training (taken from different images, see **METHODS**), and 69.9% accuracy on the training sets (**Supplementary Table S1**). For **oset-2**, the classifier separates the two categories in the test sets with an average accuracy of 68.5%, with average accuracy of 70.7% on the training sets (**Supplementary Table S2**). Similar results were obtained when the training and test sets were both sampled uniformly (with replacement) from all images (see **METHODS**). In this analysis, the classifier trained on **16K** image patches predicts test set category (**4K** patches) with accuracy of 68.7% for **oset-1**, and 69.6% for **oset-2**.

**Fig. 3** (*left + center*) illustrates examples of image patches near the 10^th^, 50^th^, and 90^th^ percentiles when the 4000 images are sorted by each contrast measure (each patch is at most 8 ranks from the exact percentile rank). **Fig. 3a** shows occlusion images from both sets, and **Fig. 3b** shows the shadow images. In our analyses, we applied our measurements to the grayscale converted (**Eq. 1**) RGB values obtained from the linearized images without any pre-normalization. This is because such operations may potentially reduce contrast cues. However, repeating these analyses where each individual linearized image patch was also normalized to lie in the range [0, 1] yielded nearly identical results (**Supplementary Fig. S2**).

**Figure 3:**
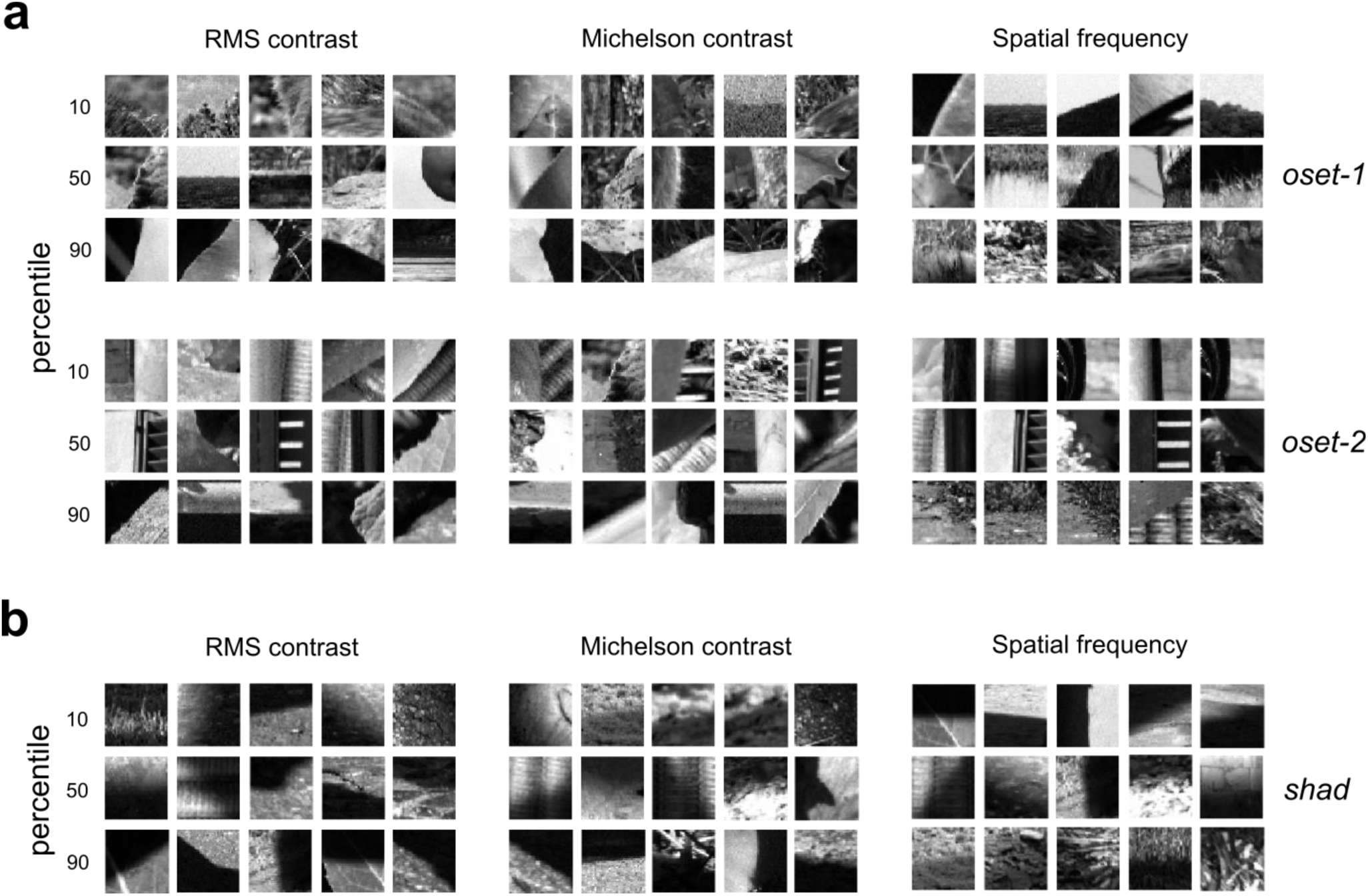
Example occlusion and shadow edges at varying percentiles of our contrast and spatial frequency measurements. Each image patch is within 8 ranks of the indicated percentile (out of N = 2000 total image patches analyzed). (a) Occlusion edges at varying percentiles of each contrast measure (RMS: *left*; Michelson: *center*) and our spatial frequency measure (𝜋_ℎ_: *right*) for both sets of occlusions (**oset-1**: *top*, **oset-2**: *bottom*). (b) Same as (a), but for shadow edges from our database.

#### Spatial frequency statistics

As we see from **Fig. 1c** and **Fig. 3**, another potential cue for distinguishing shadow edges from occlusions is that many shadows exhibit a gradual transition between luminance levels due to penumbral blur (**Casati & Cavanagh, 2019**). Therefore, we should expect that edges arising from shadows may have less energy in higher spatial frequency (SF) ranges than those arising from occlusions. Furthermore, occlusions may exist without any change in luminance, and have high-contrast high-SF textures on both sides of the boundary, whereas with shadows the high-SF texture on the darker half will generally have lower contrast. Analysis of the proportion 𝜋_ℎ_of stimulus energy at higher spatial frequencies (which we define as those frequencies above 10 cycles/image, with 20 cycles/image being the Nyquist limit) reveals that indeed there are significant differences between occlusions and shadows (**Fig. 2c**). Comparison of 𝜋_ℎ_ distribution medians (rank-sum test) yielded significant results for both **oset-1** (occlusion median = 0.076, shadow median = 0.060, *p* < 0.001) and **oset-2** (occlusion median = 0.078, shadow median = 0.060, *p* < 0.001). Similar results were obtained when we increased our cutoff for high spatial frequencies to 15 cycles/image. Averaging the amplitude spectrum obtained from each patch (**Fig 2d**) reveals more energy at higher spatial frequencies for the occlusion images compared to shadow images. **Fig. 2e, f** shows scatterplots of 𝜋_ℎ_ versus 𝑐_𝑀_and 𝑐_𝑅𝑀𝑆_ for both **oset-1** (*left*) **and oset-2** (*right*). Significant negative correlations were observed between both contrast measures and 𝜋_ℎ_ for both occlusion sets and the shadow set. That is, higher contrast edges had a smaller proportion of their energy in high spatial frequencies. This makes sense because the highest contrast edges (of either category) tend to have a strong difference in illumination over the spatial scale of the entire patch, which would give rise to a large low-spatial frequency component, as well as decreased energy at high SFs on the darker side. For occlusion **oset-1**, we obtain correlations between 𝜋_ℎ_and 𝑐_𝑀_, 𝑐_𝑅𝑀𝑆_of -0.492, -0.261, respectively (*p* < 0.001 both tests). For occlusion **oset-2**, we a obtain correlations between 𝜋_ℎ_ and 𝑐_𝑀_, 𝑐_𝑅𝑀𝑆_of -0.315, -0.145, respectively (*p* < 0.001 both tests). For the **shad** set, we obtain correlations between 𝜋_ℎ_ and 𝑐_𝑀_, 𝑐_𝑅𝑀𝑆_ of -0.414 (*p* < 0.001) and -0.052 (*p* = 0.002).

**Fig. 3a** (*right*) shows edges from each set corresponding to the 10^th^, 50^th^, and 90^th^ percentiles of 𝜋_ℎ_, and **Fig. 3b** (*right*) shows shadows at these same percentiles. We see occlusions with high values of 𝜋_ℎ_ tend to arise when one or both adjacent surfaces is comprised of high-spatial frequency textures, and that shadows with high values of 𝜋_ℎ_occur when the shadow overlays a high-spatial frequency texture pattern. Although we observe significant differences in our measure of high spatial frequency content between edges and shadows (**Fig. 2c**), we find that when we add this variable to our logistic regression classifier, we do not observe an improvement in discrimination performance. For **oset-1**, a classifier using all three cues can separate occlusions and shadows with an average performance of 66.4% accuracy on the test image sets (non-overlapping with training sets), and 69.7% accuracy on the training sets (**Supplementary Table S1**). For **oset-2**, the classifier separates the two categories in the test sets with an average accuracy of 65.9%, with average accuracy of 70.5% on the training sets (**Supplementary Table S2**). Similar results were obtained with a second analysis in which both training and test sets were sampled uniformly from the full database, with performance of 68.5% on the test set for **oset-1** and 69.4% on the test set for **oset-2**.

### Image-computable machine learning classifiers

#### Gabor filter classifier

Our analysis of image patch statistics demonstrated that simple features like contrast and spatial frequency could separate the two categories well above chance at a resolution of 40×40 pixels. However, this simple set of intuitively defined features does not exhaust all the information potentially available to human or machine observers to separate these two categories. Therefore, to gain additional insight we considered the performance of two biologically plausible image-computable machine learning classifiers.

The initial representation of local image regions in the visual cortex is computed by the population of V1 simple cells, which are well modeled as bank of multi-scale linear Gabor filters with half-wave rectified outputs. We performed binomial logistic regression on the rectified and down-sampled (MAX pooling) outputs of a bank of physiologically plausible oriented Gabor filters (**Kovesi, 2000; Field, 1987**) defined at multiple spatial scales (8×8, 16×16, 32×32). We refer to this model illustrated in **Fig. 4a** as the **GFB** (Gabor Filter Bank) classifier, and Gabor filters at one spatial scale are illustrated in **Supplementary Fig. S3**.

**Figure 4:**
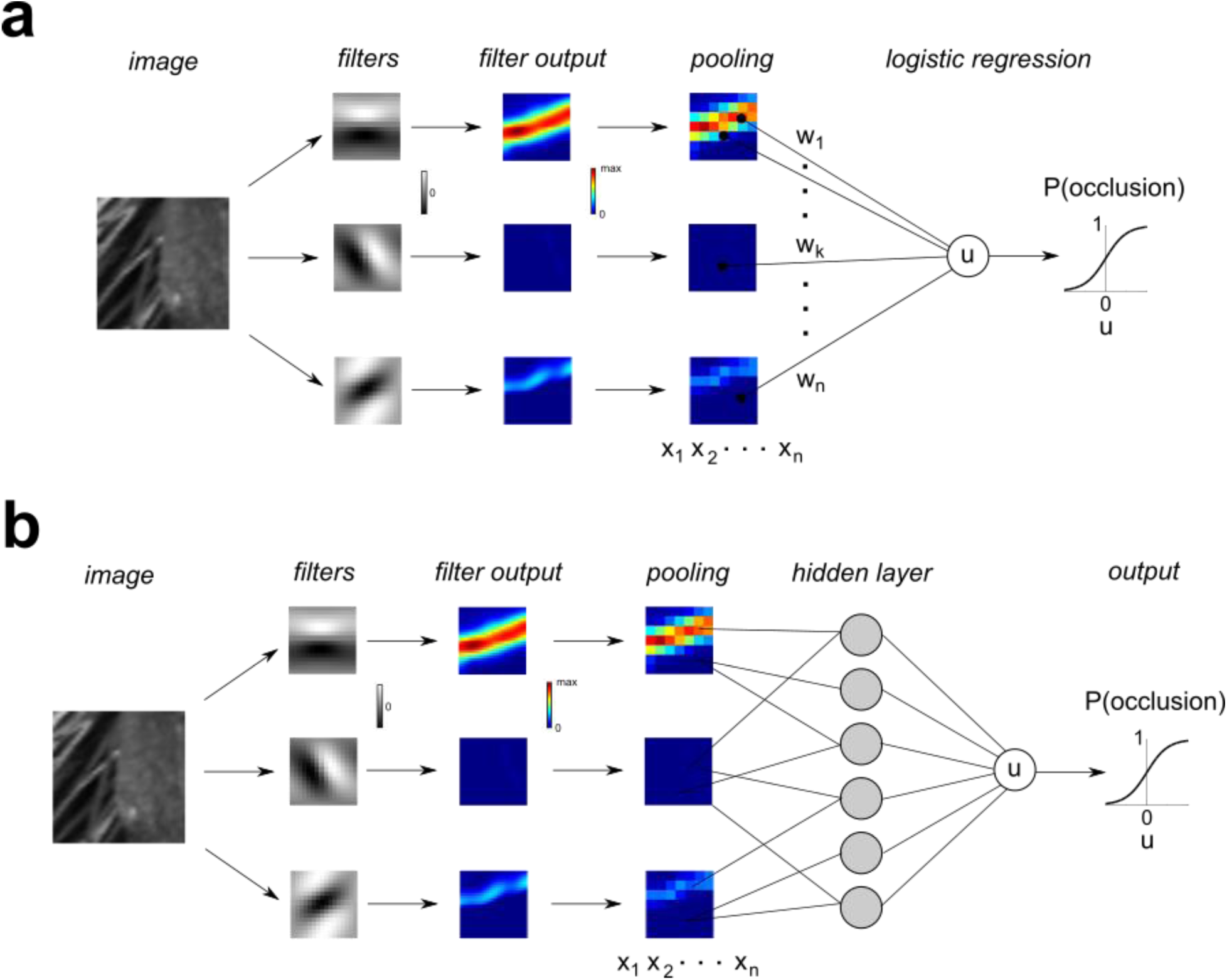
Schematic illustration of Gabor Filter Bank (GFB) and Filter-Rectify-Filter (FRF) classifier models. (a) **GFB** model. An image patch is analyzed by a bank of 72 oriented Gabor filters covering 3 spatial scales (8, 16, 32 pixels). The rectified filter outputs are down-sampled and MAX pooled to obtain a set of regressors 𝑥_1_, …, 𝑥_𝑛_. Binomial logistic regression is performed to find a set of weights that optimally predict the probability that the patch is an occlusion. b) **FRF** model. This model is identical to the **GFB** model, except the regressors 𝑥_1_, …, 𝑥_𝑛_ are then passed into a three-layer neural network having *K* hidden units (*gray circles*) to determine the probability an image patch is an occlusion edge.

The **GFB** classifier performed significantly better than the classifier defined on contrast statistics (**Fig. 2**), and its performance was robust over a wide range of regularization hyper-parameters (**Supplementary Fig. S4**). Breaking the images into 5 folds of non-overlapping images revealed average test set performance of 78.5% correct for **oset-1** (**Supplementary Table S3**), although average test set performance was slightly worse for **oset-2** (70.8% correct, **Supplementary Table S4**). Average training set performance was about the same for both sets (**oset-1**: 84.7%, **oset-2**: 82.8%). This poorer generalization ability for **oset-2** may have resulted from the relatively small size of this occlusion set (only 20 images), leading to over-fitting of the training data. When both test sets and training sets were sampled from the entire image database uniformly (with replacement), we found much better model performance on the test sets (**oset-1:** 79.9%, **oset-2:** 80.4%), most likely because they were being trained and tested on similar (or even identical) image patches (**Supplementary Table S5, Supplementary Fig. S5**).

To determine the stimulus which would be judged by the **GFB** classifier to be most likely to be classified as an occlusion (or shadow), we numerically found the stimulus to maximize (or minimize) the model output. Unfortunately, neither of these stimuli obtained via optimization visually resembles the actual image patches from either category (**Supplementary Fig. S6a**). However, we do see that the stimulus most likely to be classified as an occlusion has a sharp border, whereas the stimulus most likely to be a shadow is relatively “flat” in the middle. Therefore, we can gain better intuition for what each model is looking for by plotting the edges (of both categories) most likely to be classified as occlusions or shadows, as shown in **Fig. 5a**. Each quadrant shows a combination of actual category (vertical dimension) and predicted category (horizontal dimension). We see that the occlusions most likely to be correctly classified as occlusions (*top left*) have low contrast with a sharp boundary between regions and quite often differences in texture on opposite sides of the border. The occlusions most likely to be mis-classified as shadows (*top right*) tend to have higher contrast and a somewhat more blurred edge. The shadows mis-classified as occlusions (*bottom left*) tend to have lower contrast and/or a sharper edge. And finally, the shadows which are most likely to be correctly classified (*lower right*) have high contrast and blurry edges.

**Figure 5:**
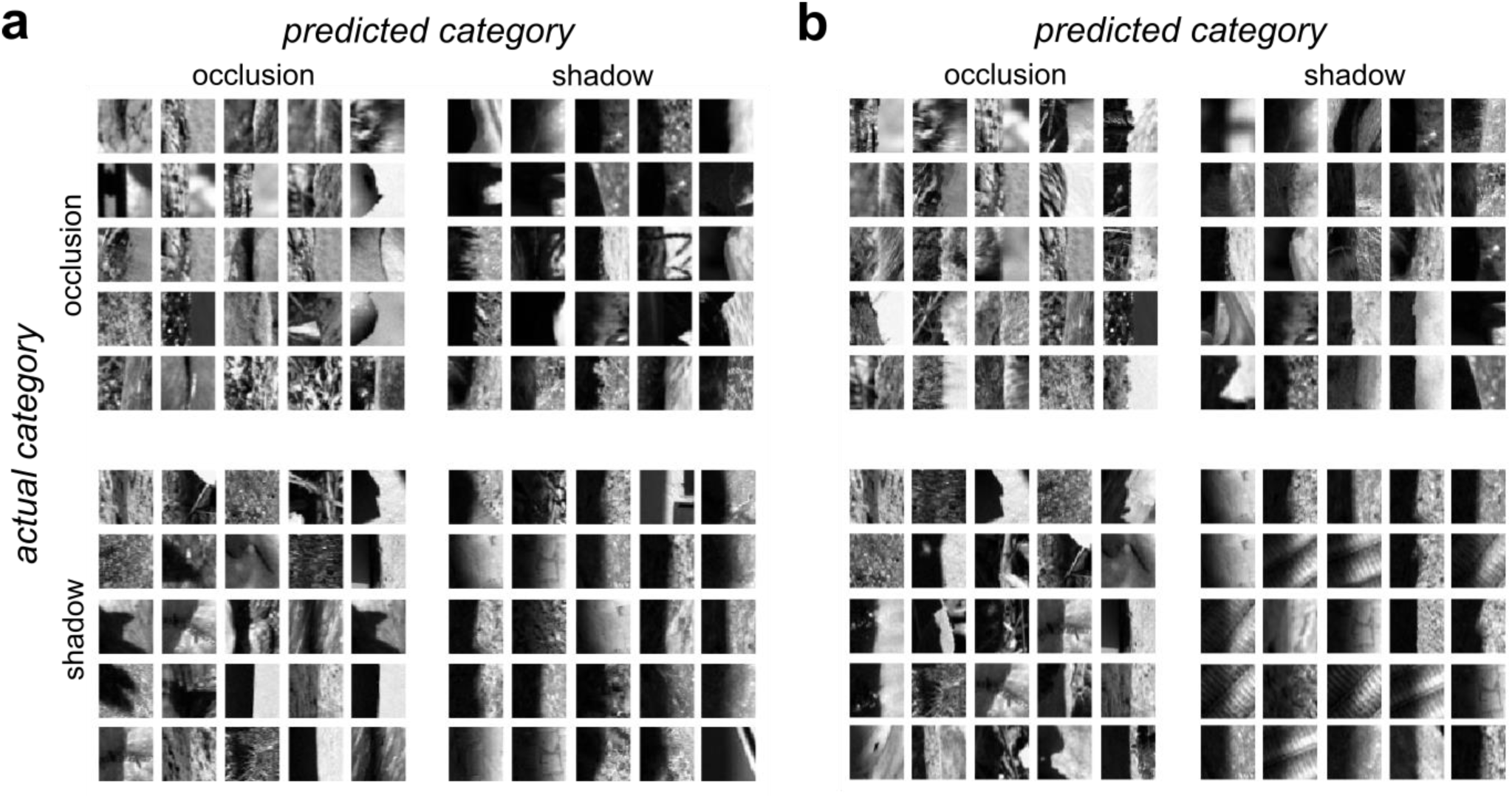
Occlusion and shadow edges with the 25 highest predicted of occlusion and shadow category membership. Actual category is indicated by the row, predicted category by the column. The upper left and lower right quadrants indicate correct classifications by the model. (a) **GFB** model (b) **FRF** model

#### Filter-rectify-filter classifier

We next considered the performance of a Filter-Rectify-Filter (**FRF**) model to classify occlusion and shadow edges (**Fig. 4b**). **FRF** models have been long used for modeling texture segmentation (**Malik & Perona, 1990; Bergen & Landy, 1991; Baker, 1999; Landy & Graham, 2004; Zavitz & Baker, 2013, 2014; DiMattina & Baker, 2019, 2021; DiMattina, 2022**) and are mathematically similar to three-layer connectionist neural networks (**Rumelhart, Hinton, & Williams 1986; Bishop, 2006; Goodfellow, Bengio, & Courville, 2016**). The first filtering layer of this network was comprised of the fixed, multi-scale Gabor filter bank used in the **GFB** classifier (**Fig. 4a**). The rectified, down-sampled outputs of this filter bank were then fed to a population of *K* hidden units (*K* = 4, 6, 10), the rectified outputs of which were linearly combined form a decision variable, which is passed into a sigmoid function to predict the occlusion probability.

When training and test sets were derived from non-overlapping sets of images, adding this hidden layer greatly improved classification performance on training sets, but yielded a more modest improvement for the test sets. For the **FRF** model with 6 hidden units (**FRF-6**), the average performance on training sets approached 90% accuracy (**oset-1**:86.3%, **oset-2**: 89.0%), but the average test set performance (**oset-1**: 80.3%, **oset-2**: 73.8%, **Supplementary Tables S6, S7**) was only somewhat better than the **GFB** model (**GFB - oset-1**: 78.5%, **oset-2**: 70.8%). Similar results were obtained for **FRF-4** (**oset-1** – *train*: 85.2%, *test*: 79.9%; **oset-2** – *train*: 80.5%, *test*: 73.5%; **Supplementary Table S8**, **S9**) and **FRF-10** (**oset-1** – *train*: 87.4%, *test*: 80.5%; **oset-2** – *train*: 91.2%, *test*: 73.6%; **Supplementary Table S10**, **S11**). As with the other models, performance was relatively insensitive to the value of the regularization hyper-parameter (**Supplementary Figs. S7- S9**). By contrast, when both the test and training sets were uniformly sampled (with replacement) from all images, for each **FRF** model and each set of stimuli (**oset-1**, **oset-2**), performance on the test sets was commensurate with that on the training sets, approaching 90% (**Supplementary Table S5**). This is because in this case the model was being tested on patches which were very similar (or identical) to those it was trained on.

As with the GFB, numerical maximization (or minimization) of FRF model output did not yield stimuli that visually resembled either category (**Supplementary Fig. S6b-d**). As before, we find the stimulus most likely to be an occlusion has a sharp boundary, whereas a shadow has a gradual boundary. Numerically finding the optimal stimuli for the hidden units in each model revealed that many hidden units seemed to be sensitive to differences in texture on opposite sides of the boundary (**Supplementary Fig. S10**). This is consistent with the idea that texture differences may be important for distinguishing shadows from surface boundaries (**Jiang et al., 2010; DiMattina 2022**).

**Fig. 5b** shows the actual and predicted categories of the image patches in **oset-1** the **FRF- 6** model predicts to have the highest probability of category membership. We see that as with the **GFB** model, occlusions correctly classified as such (*top left*) tend to have low contrast and sharp borders, and shadows correctly classified as such (*bottom right*) tend to have high contrast and blurred borders. Occlusion patches mis-classified as shadows tend to have high contrast and smoother boundaries (*top right*), whereas shadows mis-classified as occlusions (*bottom left*) have lower contrast and sharper boundaries.

### Models are sensitive to penumbral blur and texture

#### Tuning for contrast and penumbral blur

In order to understand what image features are being exploited by the **GFB** and **FRF** models to distinguish the image categories, we considered the response of the models to luminance step edges with varying levels of contrast and penumbral blur (**Fig. 6a**). **Fig. 6b** shows the response of the **GFB** model to step edges with varying degrees of penumbral blur. Different curves denote different levels of Michelson contrast (𝑐_𝑀_), which is held constant (via numerical image re-scaling) for each stimulus on the curve. We see that increasing the level of penumbral blur decreases the probability that the stimulus is classified as an occlusion edge, or equivalently makes it more likely to be classified as a shadow. Furthermore, we find that as contrast increases, the edge probability also decreases, consistent with our analyses shown in **Fig. 2**. Similar results were obtained for the **FRF** model, as illustrated in **Fig. 6c**. For both models, we see in **Fig. 6** that the sensitivity to both the blur and contrast parameters can be quite dramatic, greatly affecting the edge classification probability.

**Figure 6:**
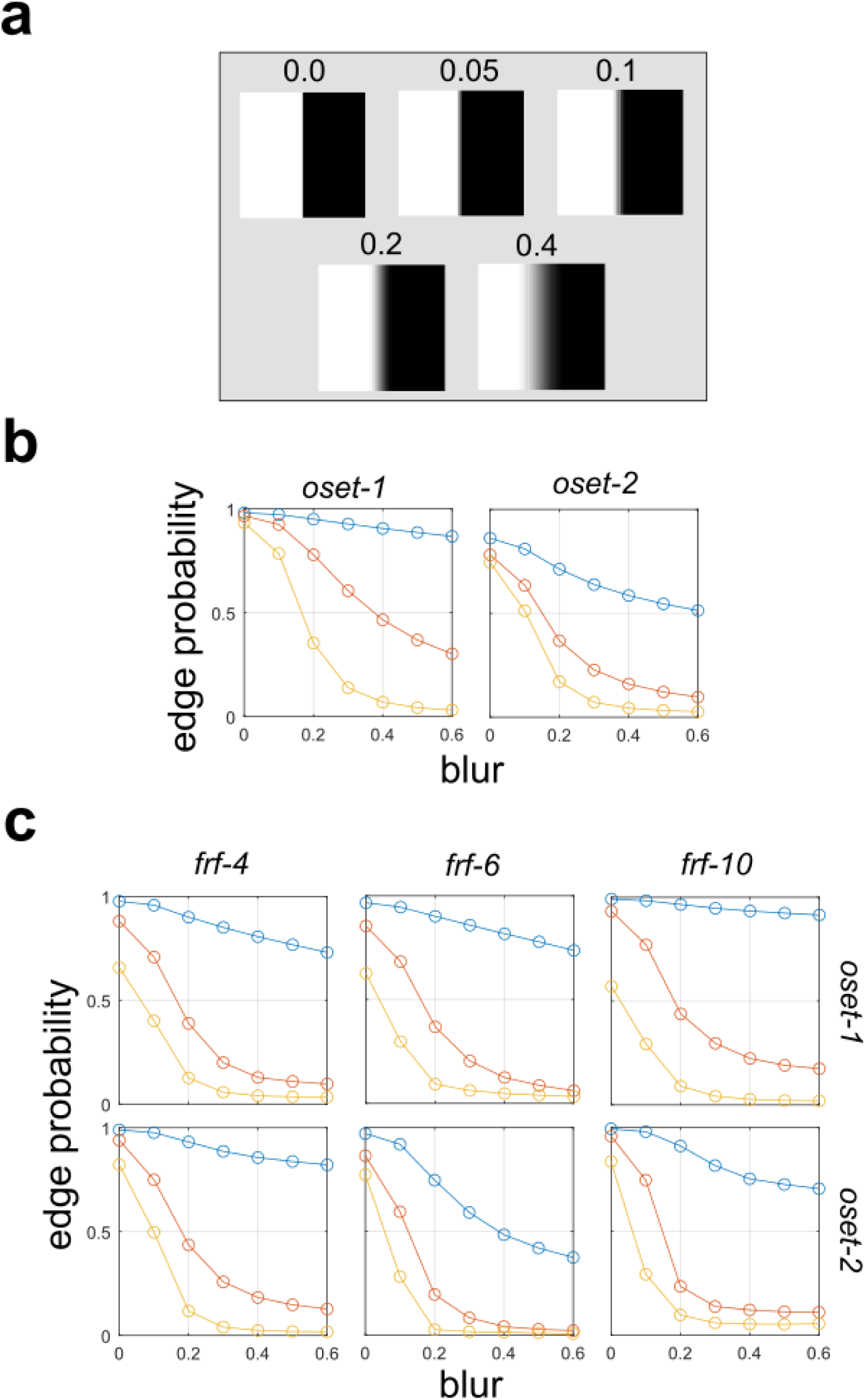
Effects of edge blur on model predictions of edge probability. (a) Test image patches with varying levels of blur. (b) Gabor Filter Bank (**GFB**) model (see **Fig. 4a**). (c) Same as (a) but for Filter-Rectify-Filter (**FRF)** models having varying numbers of hidden units (**Fig 4b**).

#### Removal of texture information

Our analyses of the two models suggests that they may be sensitive to texture differences on opposite sides of the image patch. To test the importance of texture cues more directly, we performed a simple image manipulation procedure to remove texture information from occlusions (**DiMattina et al. 2012**). Our method simply sets the intensity of each pixel on each side of the boundary equal to the mean value measured from that side. This manipulation leaves the image Michelson contrast (**Eq. 3**) unchanged since the mean intensity on each side remains identical. Examples of occlusion patches manipulated in this manner are shown in **Fig. 7a**. The output of the **GFB** and **FRF** models is an occlusion probability 𝑝 (with shadow probability being 1 − 𝑝). The output of each model is computed from the log-odds ratio 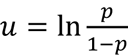 by applying the logistic link function. Therefore, we may recover the log-odds ratio by applying the inverse logistic link function σ^−1^ to the predicted occlusion probability. A positive log-odds ratio indicates that an image patch is more likely to be an occlusion, and a negative log-odds ratio indicates a shadow. **Fig. 7b** plots the log-odds ratio for both models (**FRF**: blue, **GFB**: green) for the original occlusion images (*horizontal axis*) and the texture removed images (*vertical axis*) from **oset-1** (*top*) and **oset- 2** (*bottom*). Values greater than zero indicate patches that are (correctly) classified as being occlusions, less than zero indicates patches mis-classified as shadows. We see that patches predicted to have a higher probability of being occlusions experience a much greater reduction in occlusion probability when texture is removed. This change in probability is shown explicitly in **Fig. 7c**, which plots the log-odds ratio of the original images against the difference Δ in log-odds ratios. A strong positive correlation of the original log-odds ratio with Δ was observed for each model for both **oset-1** (**FRF**: *r* = 0.9594, *p* < 0.001, **GFB**: *r* = 0.9206, *p* < 0.001, N = 191) and **oset-2** (**FRF**: *r* = 0.9755, *p* < 0.001, **GFB**: *r* = 0.9776, *p* < 0.001, N = 596).

**Figure 7:**
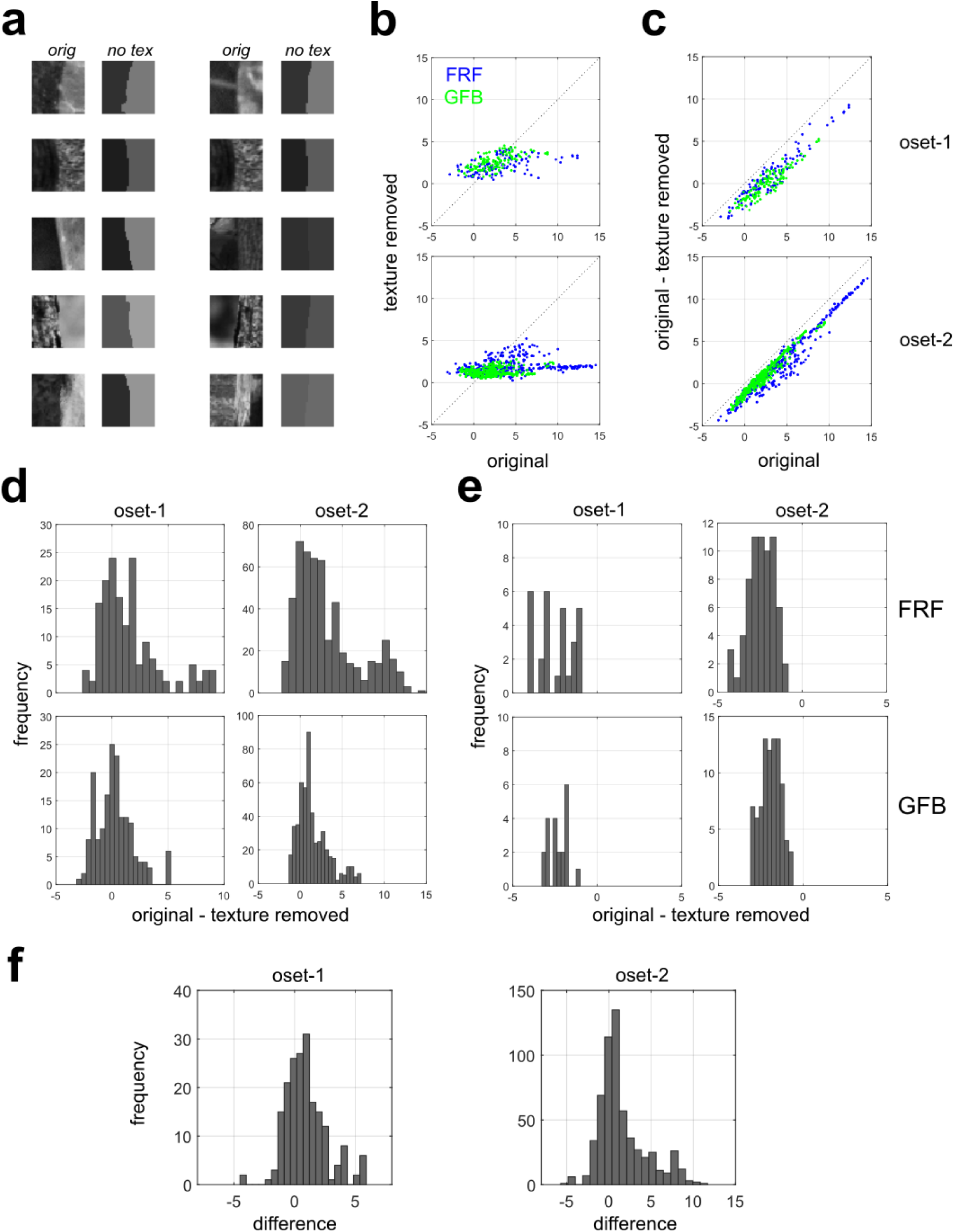
Effects of removing texture information from image patches on responses of the GFB and FRF models. (a) Averaging the pixel intensities on each side of the occlusion boundary remove texture information while leaving Michelson contrast unchanged. (b) Log-odds ratio of being an occlusion for the original images (*horizontal axis*) versus the texture-removed images (*vertical axis*). We see that images for which the model predicts a high occlusion probability experience a drastic decrease in occlusion probability when texture is removed. *Top:* **oset-1**, *Bottom:* **oset-2**. (c) Log-odds ratio of being an occlusion for the original images (horizontal axis) versus the change in occlusion probability when texture is removed (vertical axis). *Top:* **oset-1**, *Bottom:* **oset-2**. (d) Change in occlusion probability for patches correctly classified as occlusions (*Top:* **FRF**, *Bottom:* **GFB**). Positive numbers indicate a decrease in occlusion probability, negative numbers an increase in occlusion probability. *Left:* **oset-1**, *Right:* **oset-2**. (e) Same as (**d**) but for patches incorrectly classified as shadows. (f) Difference D = Δ_FRF_ – Δ_GFB_ in the reduction in edge probability with texture removal for each image. Positive values of D indicate a greater reduction in edge probability with texture removal for the **FRF** model than the **GFB** model.

Interestingly, we see that occlusions which were mis-classified as shadows (original log- odds < 0) increase their occlusion probability, likely because the texture-removal operation creates a sharp boundary (**Fig. 7a**), thereby eliminating the blur which is taken as evidence for a shadow (**Fig. 6**). **Fig. 13d** explicitly plots a histogram of Δ for correctly classified occlusions for **oset-1** (*left*) and **oset-2** (*right*) for both the FRF (*top*) and GFB models (*bottom*). For the FRF model, statistical testing reveals significantly positive (Wilcoxon rank-sum test) median value of Δ for both sets (**oset-1**: 0.716, *p* < 0.001; **oset-2**: 2.07, *p* < 0.001). By contrast, for the GFB model, for **oset-1** the median value of Δ fails to reach significance (median = 0.165, *p* = 0.138), although it is significant for **oset-2** (median = 1.073, *p* < 0.001). **Fig. 7e** is the same as **Fig. 7d** but for occlusion patches which were incorrectly classified as shadows. For the **FRF** model (*top*), we find a significantly negative median value of Δ for both sets (**oset-1**: -2.33, *p* < 0.001; **oset-2**: -2.41, *p* < 0.001). For the GFB model (bottom) we also obtain significant negative values (**oset-1**: -2.334, *p* < 0.001; oset-2: -1.861, *p* < 0.001). Similar results were obtained for **FRF-4**, **FRF-10**, as summarized in **Supplementary Table S12**.

For each pair of image patches (original and texture-removed), for each model we can compute Δ (Δ_FRF_, Δ_GFB_) and then compute the difference D = Δ_FRF_ - Δ_GFB_ in order to test the hypothesis that removal of texture information has a stronger effect on the responses of the **FRF** model than the **GFB** model (D > 0). Histograms of this difference D are shown in **Fig. 7f**, and in both cases we obtain a median significantly greater than zero (**oset-1:** 0.724, *p* < 0.001; **oset-2**: 0.755, *p* < 0.001), suggesting that the **FRF** model is more sensitive to texture information.

### Psychophysical experiment

#### Individual observer performance

A single-interval forced choice experiment was performed in which human observers classified 40×40 pixel grayscale natural image patches as being shadow or occlusion edges (see **METHODS** for details). This experiment is schematically illustrated in **Fig. 8a**, and was implemented online using Qualtrics for stimulus presentation and recording responses (**Supplementary Fig. S11**). The first survey (**QT-1**) was considered a “practice” session, although we found similar results with the “test” surveys (**QT-2**, **QT-3**) and hence included it in our final analysis. **Fig. 8b** (*left*) shows the performance (proportion correct 𝑝_𝑐_) of all N = 18 observers who completed all three online experiments (with above-chance performance on the final two). We find the median observer (*large blue circles*) had performance of 𝑝_𝑐_ = 0.69 (**QT-1**), 𝑝_𝑐_= 0.688 (**QT-2**), 𝑝_𝑐_= 0.740 (**QT-3**), with the best observer attaining performances (*black diamonds*) of 𝑝_𝑐_= 0.805, 𝑝_𝑐_= 0.755, 𝑝_𝑐_= 0.80 respectively. Performance was not significantly correlated across surveys for 2 of 3 possible pair- wise comparisons (**QT 1/2**: *r* = 0.222, *p* = 0.376; **QT 1/3**: *r* = 0.54, *p* = 0.021; **QT 2/3**: *r* = 0.129, *p* = 0.611). Therefore, for many analyses we pooled data across surveys. In these cases, the units of analysis will be referred to as *observer-surveys*. **Fig. 8b** (*right*) also shows performance quantified as 𝑑^′^ for all experiments, and we observe a strong positive correlation between 𝑑^′^ and 𝑝_𝑐_ for each survey (**QT-1**: *r* = 0.998, *p* < 0.001; **QT-2**: *r* = 0.993, *p* < 0.001; **QT-3**: *r* = 0.957, *p* < 0.001) and across all N = 54 observer-surveys (*r* = 0.985, *p* < 0.001, N = 54).

**Figure 8:**
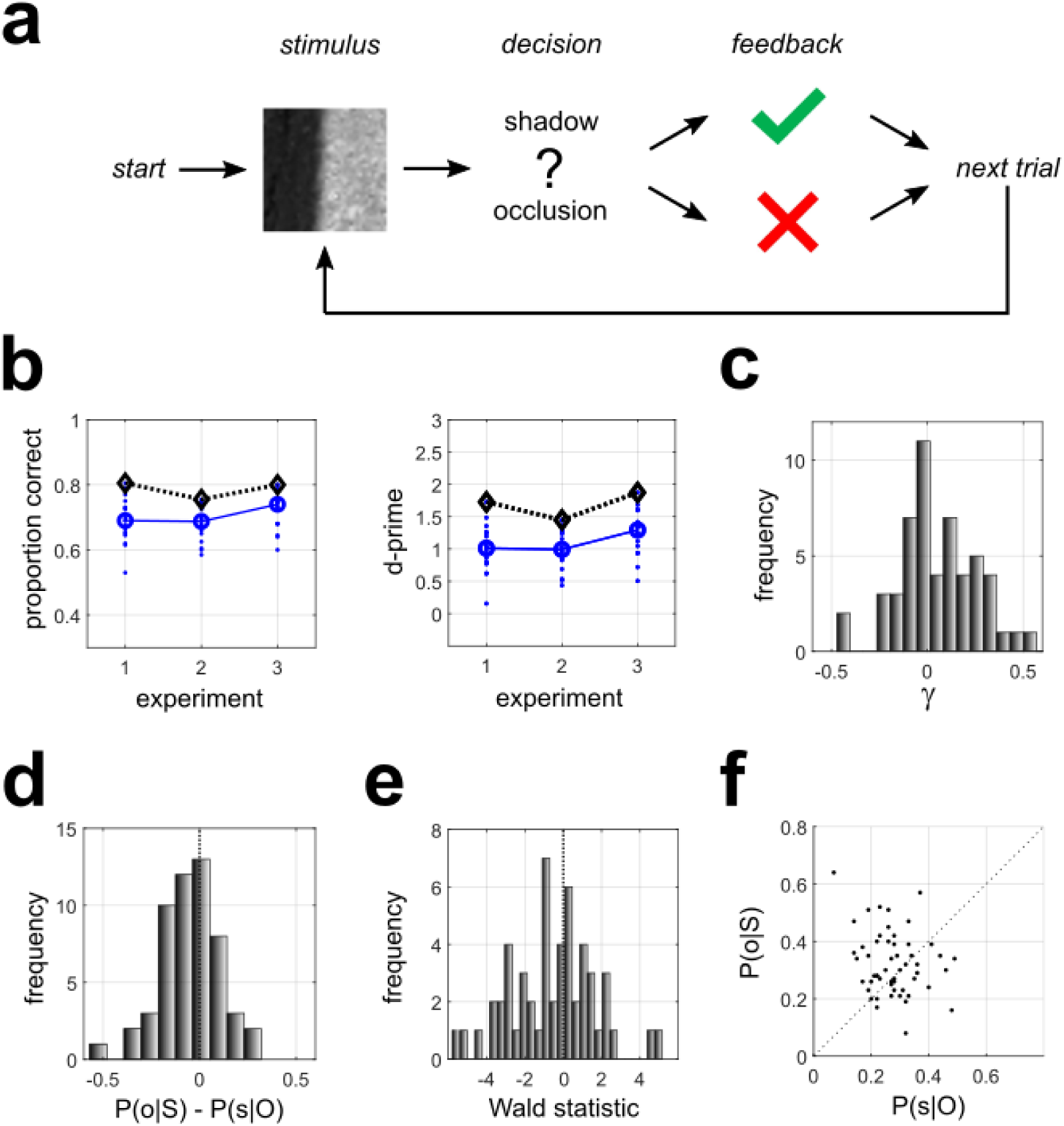
Psychophysical experiment results. (a) Schematic outline of our psychophysical task. (b) *Left:* Performance of all observers on each survey, as measured by proportion correct. Blue dots indicate individual observers. Blue circles indicate the median, and black diamonds indicate maximum performance. *Right:* Same as left panel, but performance measured as d-prime. (c) Distribution of the SDT decision criterion (γ) for all observers and surveys. (d) Distribution of the difference 𝑃(𝑜|𝑆) − 𝑃(𝑠|𝑂) in classification error probabilities. 𝑃(𝑜|𝑆) indicates the probability a shadow is mis-classified as an occlusion, and 𝑃(𝑠|𝑂) is the probability of an occlusion being mis-classified as a shadow. (e) Distribution of the Wald statistic for the binomial proportion test testing whether there is a bias in mis-classifications, with the null hypothesis of equal mis-classification probabilities. (f) Scatter plot of classification error probabilities.

Observer performance in standard signal detection theory is determined by the separation 𝑑^′^ of the distributions of the decision variables corresponding to each category, as well as the decision criterion 𝛾 applied by the observer. In our SDT analysis, 𝛾 < 0 corresponds to a bias towards judging stimuli to be occlusions, and 𝛾 > 0 corresponds to a bias towards judging stimuli to be shadows. For 2 of the 3 surveys, the median value of 𝛾 was not found to be significantly different from zero using a Wilcoxon test (**QT-1**: median = 0.097, *p* = 0.042; **QT-2**: median = 0.082, *p* = 0.119; **QT-3**: median = -0.008, *p* = 0.856). Distributions of 𝛾 for each survey are shown in **Supplementary Fig. S12**. Pooling 𝛾 across surveys (**Fig. 8c**) reveals a median which was not significantly different from zero (Wilcoxon test, N = 54, median = 0.066, *p* = 0.059). Computing the ratio |𝑑^′^/𝛾| for 𝛾 ≠ 0 (N = 51) reveals a median value of 8.78, with quartiles Q_1_ = 3.92 and Q_3_ =15.06. Therefore, any bias in the decision criterion is for most observers negligible compared to the distribution separation.

Since one of our goals is to compare human performance to that of machine observers, we want to not only compare the probability of correct response, but also the pattern of classification errors. For each observer and experiment, we measured the probability 𝑃(𝑠|𝑂) that an occlusion was misclassified as a shadow, and the probability 𝑃(𝑜|𝑆) that a shadow was misclassified as an occlusion. If there is no bias in either direction, the difference 𝑃(𝑠|𝑂) − 𝑃(𝑜|𝑆) (**Fig. 8d**) should be zero. A positive value would indicate a greater likelihood of misclassifying occlusions as shadows than misclassifying shadows as occlusions. A negative value would indicate a greater likelihood of mis-classifying shadows as occlusions than the other way around. On an individual basis, this can be assessed using a difference of binomial proportion test (*Wald test*). Doing so revealed that over 54 observer-surveys (18 observers taking 3 surveys each), in 22/54 cases a significant bias was observed towards misclassifying one or the other stimulus categories. Of these, 6/22 observer-surveys exhibited a bias to misclassify occlusions as shadows, whereas in 16/22 observer-surveys we saw a bias to mis-classify shadows as occlusions. The proportion of observer- surveys which exhibited a significant bias to mis-classify shadows as occlusions does not differ significantly from chance (binomial proportion test, N = 22, *p* = 0.055). Over the full set of observer-surveys, comparing the median Wald statistic (**Fig. 8e**) to zero (sign-rank test) failed to reach the conventional criterion for statistical significance (median = -0.675, *p* = 0.057, N = 54). **Fig. 8f** shows a scatterplot of 𝑃(𝑠|𝑂) and 𝑃(𝑜|𝑆), and we see that points are about equally likely to be above and below the diagonal.

#### Aggregate observer performance across images

In addition to analyzing performance of each individual observer, for each individual image (200 per set, for 3 image sets), we characterized the proportion of observers that classified it as a shadow or an occlusion. Doing this allows us to generate an *aggregate observer* (**AO**) model, which provides a model of average human performance against which the performance of machine classifiers (and other human observers not used to define the model) can be compared. Mathematically, the aggregate observer for K participants making a binary behavioral classification (𝑏 = 0, 1) of 𝑁 images is given by 𝒜 = {𝜋_𝑖_}^𝑛^, where 𝜋_𝑖_ indicates the proportion of observers for which 𝑏_𝑖_ = 1. The **AO** model provides a statistical model of average human performance on the task. One can generate simulated datasets stochastically using the **AO** by generating response 𝑏_𝑖_ = 1 to stimulus 𝑖 with probability 𝜋_𝑖_, and response 𝑏_𝑖_ = 0 with probability 1 − 𝜋_𝑖_. Likewise, one can generate a stimulated dataset deterministically by setting 𝑏_𝑖_ = 1 if 𝜋_𝑖_ ≥ 0.5, and 𝑏_𝑖_ = 0 otherwise.

**Fig. 9** shows the probability that each stimulus is classified as an occlusion by the **AO** (blue curves), with actual stimulus categories indicated by a black (+) symbol. We see that the overwhelming majority of the stimuli with low occlusion probability are shadows, and most stimuli with high occlusion probability are occlusions, with little bias in either direction. **Fig. 9b** breaks down these plots by the actual category of the image. **Table 1** indicates the confusion matrices for each survey obtained from the **AO** using a deterministic rounding rule that sets 𝑏𝑖 = 1 if 𝜋_𝑖_ ≥ 0.5, and 𝑏_𝑖_ = 0 otherwise.

**Figure 9:**
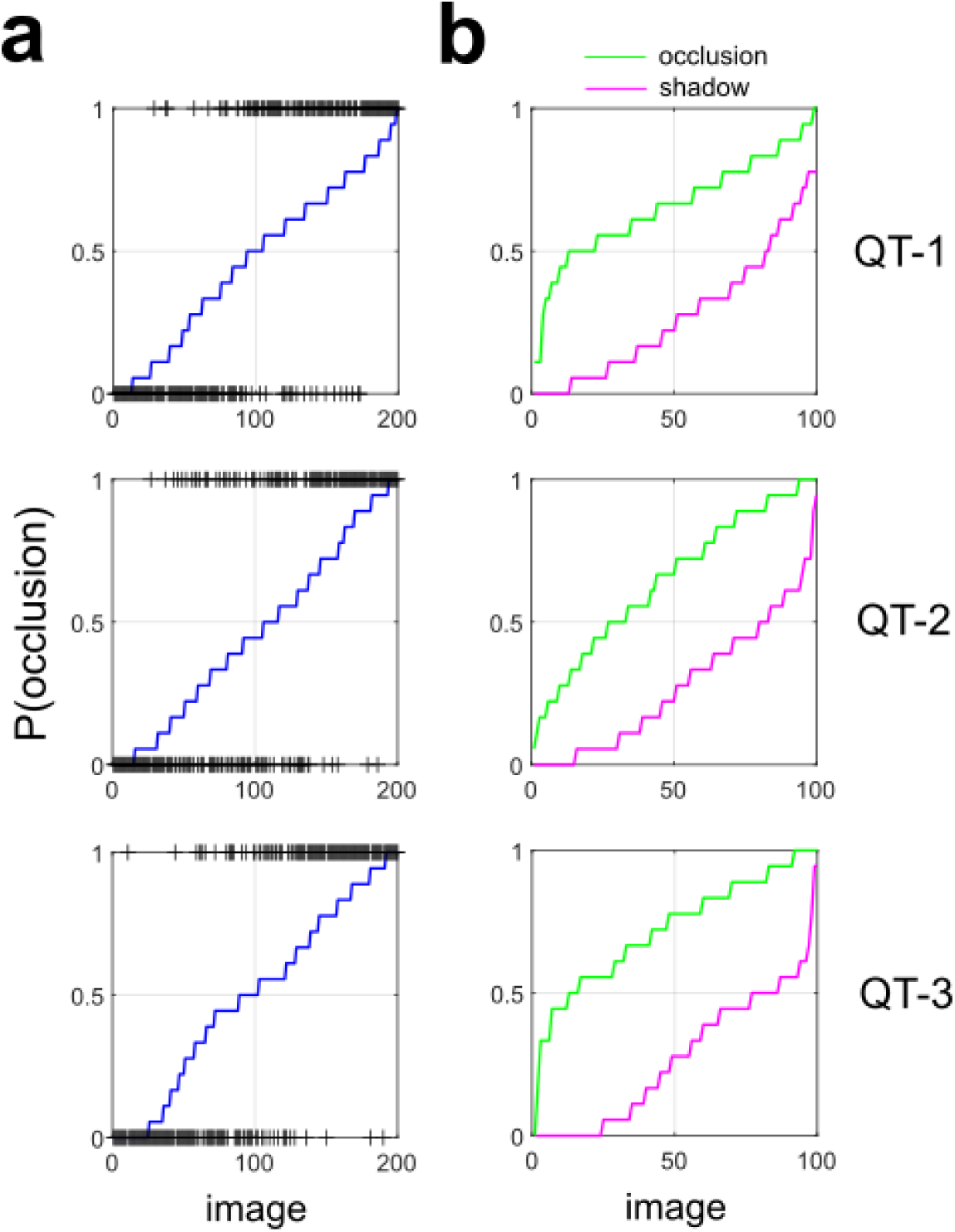
Human performance on individual images. (a) Predicted probability of being an occlusion edge by the **AO** (*blue curve*). Image patches are sorted by their probability. Actual category is indicated by a (+). (b) Occlusion probability predicted by the **AO** for each occlusion edge (*green curve*) and each shadow edge (*magenta curve*), sorted by occlusion probability.

**Table 1:** AO confusion matrix for all experiments.

|  | **QT-1** |  | **QT-2** |  | **QT-3** |  |
| --- | --- | --- | --- | --- | --- | --- |
|  | **predicted** |  | **predicted** |  | **predicted** |  |
| **actual** | *occlusion* | *shadow* | *occlusion* | *shadow* | *occlusion* | *shadow* |
| *occlusion* | 88 | 12 | 74 | 26 | 88 | 12 |
| *shadow* | 19 | 81 | 21 | 79 | 24 | 76 |

From these confusion matrices, we observe that the proportion correct for the **AO** (**QT-1**: 84.5%, **QT-2**: 76.5%, **QT-3**: 82.0%) is not significantly different from the best individual observer (95% CI of the difference: **QT-1**: [-0.034, 0.114], **QT-2**: [-0.074, 0.094], **QT-3**: [-0.057, 0.097]). Therefore, there seems to be a “wisdom of crowds” effect whereby averaging over a population of observers yields an aggregate observer that does as well as the best individuals. For each experiment, we tested whether the proportions of errors were biased in one or another direction, with the null hypothesis being that the two proportions were equal. A binomial proportion test revealed no significant difference for **QT-1** (*p* = 0.281, N = 31), **QT-2** (*p* = 0.559, N = 47), or **QT- 3** (*p* = 0.066, N = 36). Therefore, the **AO** is just as likely to mis-classify occlusions as shadows as it is to make the opposite error.

### Comparison with machine classifiers

#### Correlation between log-odds ratios

For each image, the aggregate observer model (**AO**) specifies a probability that the image is an occlusion, much in the same manner as two machine classifiers investigated in an earlier paper. In order to see if images which were more likely to be categorized as occlusions by the **AO** were also more likely to be categorized as such by various machine observers, we developed a simple analysis procedure, which we now describe. Briefly, each of these machine classifiers at its final stage computes an intermediate variable 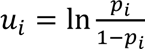 (the log-odds ratio) which is them passed through a monotonic sigmoidal non-linearity in order to compute a probability 𝑝_𝑖_ of image 𝑖 being an occlusion. By applying the inverse sigmoid function 𝑢_𝑖_ = 𝜎^−1^(𝑝_𝑖_) to the probability computed by the model, we recover the value of the log odds ratio, which unlike the occlusion probabilities computed over the set of images (**Supplementary Fig. S13**) enjoys an approximately normal distribution. Applying this same inverse sigmoid function to the occlusion probability 𝑝_𝑖_ computed from the **AO** permits recovers the corresponding log-odds 𝜎^−1^(𝑝_𝑖_) when 0 < 𝑝_𝑖_ < 1. Note that 𝜎^−1^(0) and 𝜎^−1^(1) are not well defined (being −∞ and +∞, respectively).

By correlating the log-odds ratios over the set of images, we can determine the extent various machine classifiers exhibit agreement with human observers. **Fig. 10** shows scatter-plots of the log odds ratios obtained from human and machine observers applied to all images in each survey (**QT-1**, **QT-2**, **QT-3**) for both the **GFB** (*left column*) and **FRF-10** (*right column*) model. To test whether there was a stronger correlation between the **GFB** and **FRF** model and human behavior, we computed the Spearman correlation between the log-odds ratios from the **FRF** and **GFB** model and the **AO** for N = 1000 bootstrapped samples. We made use of Spearman rank-order correlation 𝜌 instead of Pearson’s *r* because some images were classified by all observers as shadows or occlusions, so that 𝑝_𝑖_ = 0 or 1, making 𝜎^−1^(𝑝_𝑖_) is undefined. We found that for all surveys a larger Spearman correlation between the **FRF** and the **AO** (**QT-1**: 𝜌 = 0.696, **QT-2**: 𝜌 = 0.587, **QT-3**: 𝜌 = 0.775, *p* < 0.001 all tests) than the **GFB** and **AO** (**QT-1**: 𝜌 = 0.604, **QT-2**: 𝜌 = 0.568, **QT-3**: 𝜌 = 0.697), and for 2/3 surveys the 95% CI of the difference did not contain zero (**QT-1**: [0.011, 0.183], **QT-2**: [-0.067, 0.118], **QT-3**: [0.010, 0.165]). Pooling all three surveys also revealed a significantly larger correlation between the **FRF** and **AO** (𝜌 = 0.692) than the **GFB** and **AO** (𝜌 = 0.624), with a median difference Δ𝜌 = 0.068 and the 95% CI of the difference [0.022, 0.118] not containing zero. Similar results were obtained with **FRF-6**, with median difference (pooling acoss surveys) between the correlations of Δ𝜌 = 0.073 and 95% CI of the difference [0.028, 0.123] not containing zero. This analysis suggests that the **FRF** model is better than the **GFB** model at capturing whatever aspects of the image humans are utilizing.

**Figure 10:**
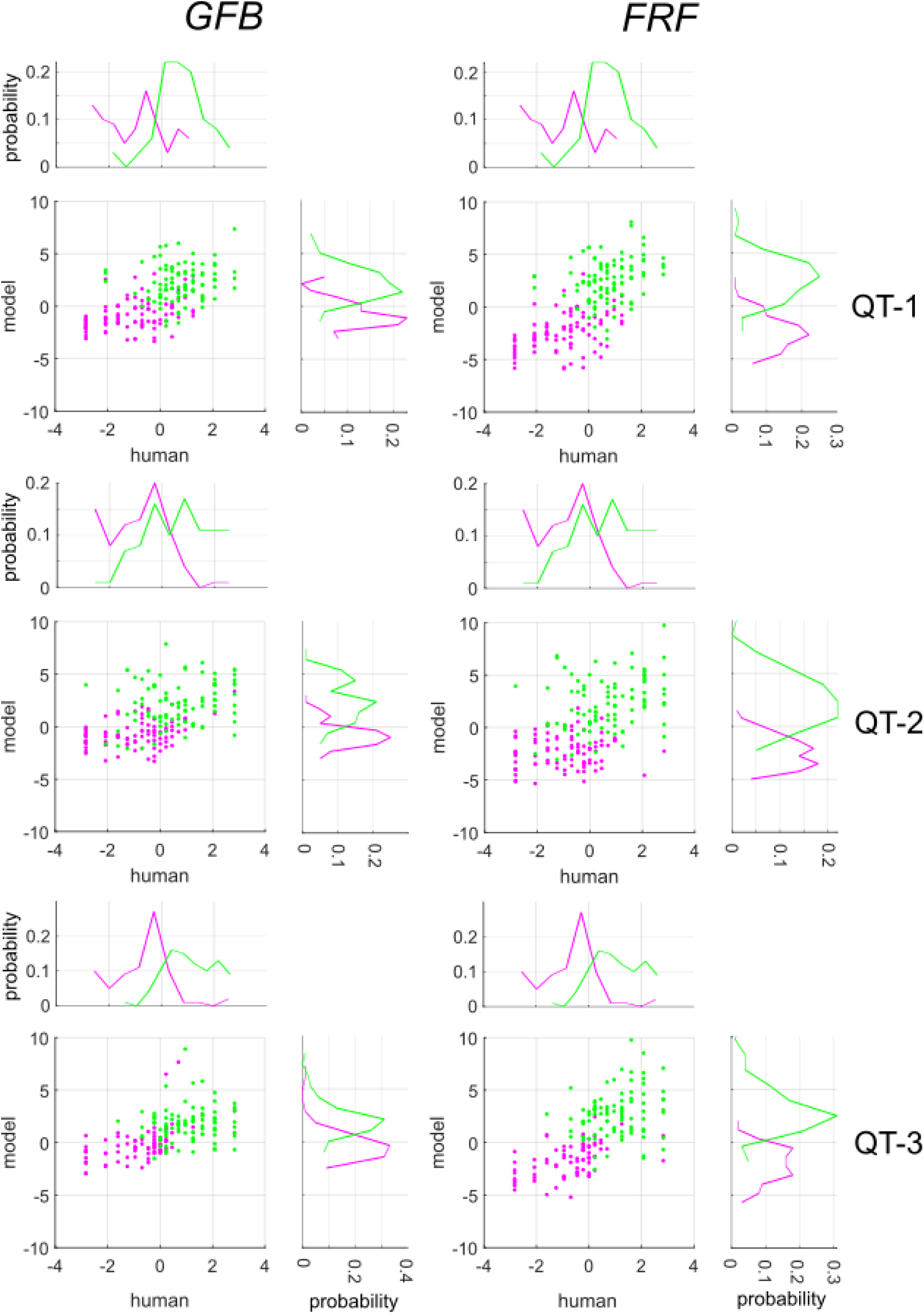
Log-odds ratios of human observers (**AO,** horizontal axes) and **GFB**, **FRF** models (vertical axes) for each image for all three surveys. Magenta symbols indicate image patches that are occlusions, green symbols indicate shadows. On the margins of each scatterplot we show the distributions of the log-odds ratios for each image category.

#### Decision variable correlation

We further explored the relative efficacy of the **FRF** and **GFB** models in accounting for human behavior using Decision Variable Correlation (**DVC**), an extension of signal detection theory (**Sebastian & Geisler, 2018**). This novel methodology assumes that for classification tasks both a human observer and a model (or two humans) make use of an internal, unobserved decision variable which each compares to some criterion to decide which stimulus was presented. Assuming the joint distribution of the human and model decision variables is a bivariate normal distribution, by analyzing the patterns of agreement and disagreement between the model and human, it is possible to recover the covariance. DVC is particularly powerful for model selection, since it is possible for two models to predict performance (as percent correct) equally well, while not agreeing on which particular stimuli the human and model classify into each category. DVC has been fruitfully applied in several domains, including texture segmentation (**DiMattina & Baker, 2019**), stimulus detection in noise (**Sebastian, Semiller, & Geisler, 2020**), speed perception (**Chin & Burge, 2000**), and avian audition (**Henry & Abrams, 2021**).

**Fig. 11a** shows for surveys **QT-1**, **QT-2**, **QT-3** the DVC between each individual observer and each of the three models: **GFB**, **FRF**, and **AO**. We see for **QT-1** a significantly larger median DVC for the **FRF** model than **GFB** model (sign-rank test, median **FRF**-**GFB** = 0.061, *p* = 0.001, N = 18), and furthermore a significantly larger median DVC for the **AO** model than either **FRF** (median **AO**-**FRF** = 0.052, *p* = 0.043) or **GFB** models (median **AO**-**GFB** = 0.114, *p* = 0.006). For **QT-2**, we do not see a significant difference in DVC between the **FRF** and **GFB** models (median **FRF**-**GFB** = -0.008, *p* = 0.349), but do see significantly better performance by the **AO** than either **FRF** (median **AO**-**FRF** = 0.22, *p* < 0.001) or **GFB** (median **AO**-**GFB** = 0.192, *p* < 0.001). Interestingly, for **QT-3** we see better median performance by the best machine classifier (**FRF**) than the **AO** which was derived from other humans (median **AO**-**FRF** = -0.146, *p* = 0.022), as well as better performance the **GFB** classifier (median **FRF**-**GFB** = 0.090, *p* = 0.02), which in turn exhibited the same performance as humans (median **AO**-**GFB** = -0.037, *p* = 0.514). **Fig. 11b** combines DVCs from all surveys. Here we observe significantly better performance for the **FRF** model than the **GFB** model (median **FRF**-**GFB** = 0.045, *p* = 0.005), although we see the best performance for the **AO** model (median **AO-GFB** = 0.142, *p* = 0.001; median **AO**-**FRF** = 0.061, *p* = 0.021). **Fig. 11c** shows histograms of these differences between the DVCs for all model comparisons, combined across the three surveys.

**Figure 11:**
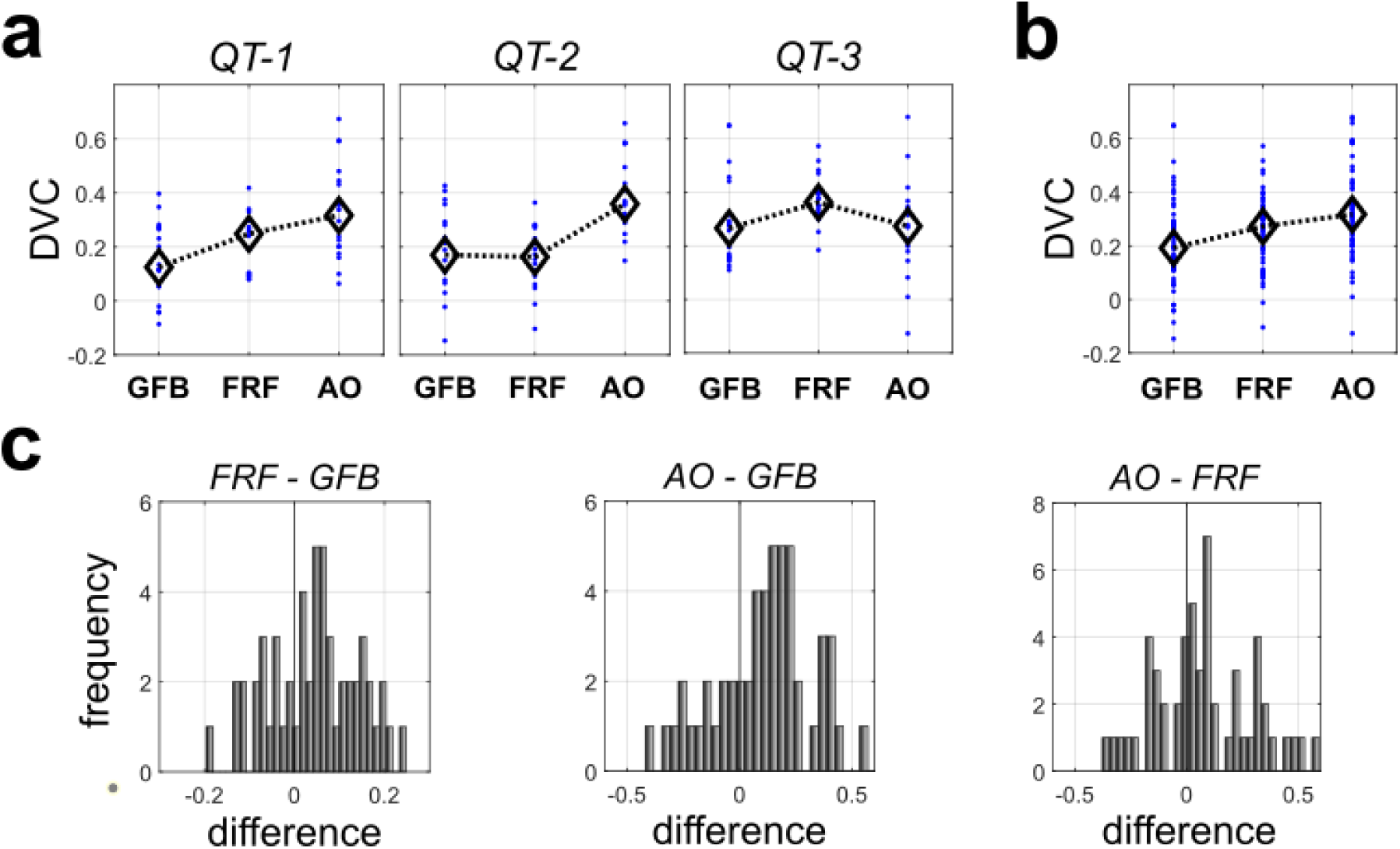
Decision-variable correlation analysis. (a) DVC between models and individual observers for each experiment. (b) Same as (a) but pooled across experiments. (c) Histograms of differences in DVC for all possible comparisons of the three models.

#### Analysis of mis-classification patterns

Another way to compare the performance of the models to human observers is to examine the patterns of classification errors. If there is no bias in either direction in the pattern of mis-classification errors, then the proportion of mis-classifications of shadows as occlusions (or vice versa) should not differ from 0.5. This can be assessed for each machine classifier (**GFB**, **FRF**) and the **AO** using a simple binomial proportion test. Here we list the proportion of classification errors which involved classifying the patch as an occlusion (*occ/all*), together with the 95% CI of the null hypothesis of equal probabilities of errors in either direction.

We see from **Table 2** that for two surveys (**QT-1**, **QT-3**) the **GFB** is biased to make errors in the direction of shadows and away from occlusions, consistent with our previous observations. By contrast, for the **FRF** model, we only observe 1 survey with a bias (**QT-2**), this time in the direction of errors in favor of occlusions. Similarly, the **AO** only shows one survey with a bias (**QT-3**). Pooling across all three surveys (**QT-1**, **2**, **3**), for the **GFB**, 35/111 mis-classifications are in the direction of occlusions, with *p* < 0.001 (binomial proportion test, N = 111). By contrast, for the **FRF** model 32/56 mis-classifications are in the direction of occlusions, and we fail to see a significant bias (*p* = 0.3496, N = 56). Similarly, for the **AO**, 50/114 mis-classifications favor occlusions, exhibiting no significant bias (*p* = 0.2234, *N* = 114). Thus, we see a much stronger agreement between the **FRF** model and average human behavior (**AO**) based on patterns of mis-classification.

**Table 2:** Number of mis-classifications of shadows as occlusions by each model.

|  | **QT-1** |  | **QT-2** |  | **QT-3** |  |
| --- | --- | --- | --- | --- | --- | --- |
|  | *occ/all* | *CI-95* | *occ/all* | *CI-95* | *occ/all* | *CI-95* |
| **GFB** | **10/33\*** | [10.9, 22.1] | 13/34 | [11.3, 22.7] | **12/44\*** | [15.5, 28.5] |
| **FRF** | 9/17 | [4.5, 12.5] | **17/23\*** | [6.8, 16.2] | 6/16 | [4.1, 11.9] |
| **AO** | 12/31 | [10.0, 21.0] | 26/47 | [16.8, 30.2] | **12/36\*** | [12.1, 23.9] |

## DISCUSSION

### Summary of contributions

Here we analyzed the statistical properties of two important classes of luminance edges: (1) occlusion edges, and (2) shadow edges. Distinguishing these two possible edge causes has important implications for understanding perceptual organization, as one kind of edge indicates a surface boundary, whereas the other indicates a change in illumination. Following previous work (**Vilankar et al., 2014; DiMattina et al., 2012**), we measured basic contrast statistics from each category, demonstrating that occlusions exhibit both the highest observed contrasts, as well as the lowest contrasts. We also measured spatial frequency properties of each image category, demonstrating more high spatial frequency content in occlusion edges than shadows. V1 neurons are known to be sensitive to both spatial frequency and contrast (**Albrecht & Hamilton, 1982; Parker & Hawken, 1988**), so relevant information for discriminating these edge categories is potentially extracted at the earliest level of cortical processing. Implementing a simple Gabor Filter Bank (**GFB**) model inspired by V1 receptive field properties, we demonstrated that at a resolution of only 40×40 pixels these two edge categories can be distinguished at nearly 80% accuracy. Furthermore, we demonstrated that this model exhibits strong tuning to penumbral blur, an important shadow cue (**Mammasian et al., 1998**; **Elder, Trithart, Pintille & MacLean, 2004; Elder & Zucker, 1998**; **Elder, 1999; Casati & Cavanagh 2019**). To investigate the performance of models which can also take texture into account, we demonstrate that a Filter-Rectify-Filter (**FRF**) model, commonly used to model second-order vision (**Baker, 1999; Landy & Graham, 2004; Zavitz & Baker, 2013, 2014; DiMattina & Baker, 2019**), can perform at greater than 80% accuracy.

In order to relate the performance of models to that of human observers, we implemented a psychophysical experiment in which observers completed the same shadow/occlusion classification task as we investigated in the models. The best human observers performed at levels commensurate with the models, although the median observer was slightly worse. Breaking down responses on an image-by-image basis, we found that while both the **FRF** and **GFB** models exhibited positive correlations with human responses, on average the **FRF** models exhibited greater correlations with human performance. Since the **FRF** model is more sensitive to texture, this is consistent with the idea that locally available texture cues may play an important role. This finding that local cues can be fruitfully exploited to assist global scene interpretation is consistent with many previous studies on natural edge detection (**McDermott, 2004; Zhou & Mel, 2008; DiMattina et al., 2012; Ing et al., 2010**), classification (**Vilankar et al., 2014; Ehinger, Adams, Graf, & Elder, 2017; Breuil et al., 2019**), and figure-ground assignment (**Burge, Fowlkes, & Banks, 2010; Ramenahalli, Mihalas, & Niebur, 2014**).

### Relationship to closely related work

Most psychophysical studies concerning shadows have investigated the essential role that shadows play in three-dimensional scene interpretation (**Ramachandran, 1988; Elder et. al. 2004; Mammasian et al., 1998; Rensink & Cavanaugh, 2004; Wilder, Adams, & Murray, 2019**). By contrast, to our knowledge, only two previous psychophysical studies directly speak to the issue of what locally available information can be used to distinguish shadows from other edge causes like occlusions (**Vilankar et al., 2014; Breuil et al., 2019**).

A previous investigation by **Vilankar et al. (2014)** characterized the statistical properties of achromatic occlusion edges and non-occlusion edges, a “catch-all” category including shadows, material changes, and surface orientation changes. Although they used local analysis windows similar in size to our own (they used rectangular windows of 81×40 whereas we used square 40×40 windows), the images which they analyzed were at high resolution (1920×2560) as opposed to the lower resolution (576×786) images we analyzed. The consequence of this is that their analysis was more “local” than our own, with their smaller dimension (40) comprising only 1.56% of the larger dimension (2560) of the image patch, whereas our 40×40 patches subtended 5.08% of the larger dimension (786). Finally, their edges were initially identified using a Canny edge detector algorithm before being classified as occlusions or non-occlusions by human observers. As they note, this process biases their samples towards edges containing stronger luminance differences, although they also analyzed a smaller set of hand-labeled edges and obtained similar results.

In their paper, they demonstrated that occlusions had higher contrast, on average, than non-occlusions. We found that when the comparison was restricted to occlusions and shadows, both the highest contrast patches as well as the lowest-contrast patches were occlusions (**Fig. 2**). Furthermore, when we analyzed occlusions from the subset of 19 images in **oset-1** that were also used in their study, the results were consistent with our general findings for both **oset-1** and **oset-2** (**Supplementary Fig. S1**). Therefore, it is unlikely that this discrepancy with our results arises from their use of a more limited image set (N = 38 images versus N = 100). It is important to note that there are two additional key differences which make direct comparison difficult: (1) We did not consider any non-occlusions apart from shadows in our analysis, and (2) They did not break down their non-occlusion stimuli into different categories in their analysis. Thus, it is possible that their finding that non-occlusions had lower contrast than occlusions would not have held if they had restricted themselves to only consider shadows.

Another interesting point of contact with **Vilankar et al. (2014)** is that in their study they plotted the luminance profiles of different categories of edges, including occlusions, non-occlusions, and shadows. They note a more gradual change in luminance for non-occlusions, including shadows, than for occlusions. All the image-computable models trained to distinguish shadows from occlusions exhibited tuning to penumbral blur, with increased blur leading to higher probability of an edge being classified as shadow (**Fig. 6**). Previous experiments suggest that humans are adept at distinguishing sharp from blurred edges (**McIlhagga & May, 2012; Ahumada & Watson, 2011**), and our analysis suggests this ability may help solve an important problem in natural scene analysis.

Another closely related study which served as a strong motivation for the present work is the study by **Breuil et al. (2019)** demonstrating the importance of chromatic cues for accurate edge classification as luminance or material. One interesting finding to arise from their study is that performance in the task for human subjects increased with edge size, whereas a similar improvement in performance was not observed with machine classifiers defined using luminance and chromatic cues. A reasonable interpretation of this result is that human observers were able to exploit textural cues which were not incorporated into the machine classifiers. Consistent with this interpretation, the present work also demonstrates that texture cues can be potentially exploited for distinguishing shadows from occlusions. Training “Filter-Rectify-Filter” (**FRF**) style classifiers revealed hidden units exhibiting sensitivity to differences in texture (**Supplementary Fig. S10**), and removing texture from the image patches lead to a drastic reduction in occlusion probability for those images most likely to be occlusions (**Fig. 7**). The utility of texture for distinguishing changes in illumination from changes in material is relatively unsurprising, as different surfaces usually have different textures. However, what is perhaps quite surprising is our finding of just how much information highly localized texture differences (40×40 pixels) can contribute.

### Limitations and future directions

The present study has several limitations, each of which provides a fruitful avenue for future work. Firstly, as in many other studies of natural perception and computer vision (**Mely et al., 2016; Zhou & Mel, 2008; Ramachandra & Mel, 2013; DiMattina et al., 2012; McDermott, 2004; Breuil et al., 2019; Martin et al., 2004**), we rely on image features annotated by human observers in a relatively small number of visual scenes. Therefore, the validity of such analyses relies on the assumption that the image sets chosen, and the region within these images which are annotated by human observers, are indeed representative of the natural visual environment.

Secondly, we focus here only on achromatic cues for shadow/occlusion classification, as previous work has rigorously investigated color and its importance for shadow edge/material edge classification (**Breuil et al., 2019**). It would be of great interest for future work to incorporate color into our neural network analyses. Given the potential utility of chromatic information for distinguishing illumination from material (**Olmos & Kingdom, 2004; Kingdom, 2019**), we might expect to find hidden units which learn to detect different colors on opposite sides of the boundary. Indeed, studies of deep neural networks trained on RGB images for classification tasks have been shown to learn filters specialized for detecting color edges while being insensitive to luminance (**Krizhevsky, Sutskever, & Hinton, 2012; McIlhagga & Mullen, 2018**). Despite the obvious interest of including chromatic cues, our work clearly demonstrates that they are by no means necessary for accurate shadow edge classification from local cues. A simple perusal of the grayscale images in **Fig. 1a** serves as proof in fact that while most certainly useful, chromaticity is by no means necessary for edge classification, and perceptual organization more broadly.

The analyses in this paper simply demonstrate that certain cues (contrast, texture, penumbral blur) are potentially useful for solving this natural vision problem, and that simple neural computations can exploit these cues. It does not tell us exactly the rules by which these cues are combined by human observers (**Jones, 2016, Kingdom et al., 2015, Ernst & Banks, 2002**), and what cues takes precedence in cases of conflict. One way this can be addressed experimentally would be to create parametric stimuli closely resembling natural stimuli, and then systematically manipulating these stimuli to determine which aspects drive behavior. For instance, our results suggest that we could create a synthetic stimulus which would be likely classified as an occlusion by a human observer by juxtaposing two different naturalistic textures with a sharp luminance border super-imposed. As we blur the border and make the textures similar or identical, observers may be more likely to classify the patch as a shadow. By varying cues simultaneously and exploring effects on stimulus categorization, we can fit and compare multiple cue combination models to better understand how texture, blur and color are integrated for shadow edge detection.

## METHODS

### Shadow and occlusion databases

#### Image sets and hand-labeling procedures

We obtained a set of N = 47 shadow images (786×576 resolution) taken from the McGill calibrated color image database (**Olmos & Kingdom, 2004**). The set of 47 images contained both natural and manmade objects. All selected images contained clear figure-ground organization and detectable shadow boundaries. Authors CD and BG with two other undergraduate researchers (LA, MD) used the image manipulation program GIMP (https://www.gimp.org/) to label the most noticeable shadows within the set of images. Each student was instructed to label the shadows individually, so their labels reflected their independent judgment. Examples of images from our shadow database (**shad**) are shown with the labeling of the shadow contours in each image by one observer (MD) shown in **Fig. 1a** (*top*). A binary union of the labels from all 4 observers is shown in **Fig. 1a** (*bottom*).

Image patches containing surface boundaries were obtained from two sets of images. One was a previously developed image database (**DiMattina et al., 2012**), also derived from a subset of N = 100 images (786 x 576) from the calibrated McGill image set. We refer to as this set of images as occlusion set-1 (**oset-1**). One potential limitation of **oset-1** is that the images in this set have almost no overlap with our set of N = 47 shadow images. Therefore, it at least possible that any statistical differences observed between shadow edges and occlusion edges could be a function of there being different surfaces present in the two image sets. Therefore, we developed a second occlusion database by having the same observers label all the occlusions in a subset of N = 20 images taken from our larger database of 47 shadow-labeled images. We refer to this second database of occlusions taken from a subset of our 47 shadow images as occlusion set-2 (**oset-2**). Two example images from this set (**oset-2**) with labeled occlusions are shown in **Fig. 1b** (*top*), together with a binary label of the occlusions from all 4 observers (**Fig. 1b**, *bottom*). Our hand-labeled shadow database and both occlusion databases are freely available to the scientific and engineering communities at https://www.fgcu.edu/faculty/cdimattina/.

#### Image patch extraction methods

Both occlusion edges and shadow edges were extracted from the image as described in **DiMattina et al. (2012)**. Briefly, random pixels labeled as edges by a single observer were chosen, cycling through the set of observers who labeled each image (CD, LA, BG, MD). To ensure patches where the boundary separates two regions of approximately equal size, with no extraneous edges, a patch centered on an edge was only accepted for inclusion if the composite edge map (pooled over all observers) for the candidate patch comprised a single, connected dividing the patch into two regions, the smaller of which comprised at least 35% of the pixels. This procedure excludes T-junctions or highly convex edges, yielding mostly smooth edges as is evident from the image patches of both categories (occlusions, shadow) shown in this paper. We focused our analyses on 40×40 image patches, which in a 786×576 image constituted roughly 5% of the larger dimension, ensuring our analysis focused on relatively local cues. Representative patches from each set are shown in **Fig. 1c**.

### Analysis of image patch statistics

#### Contrast statistics

Previous investigations have demonstrated interesting differences in the statistical properties of occlusion patches and non-occlusions (including shadows) and shown that these statistics can be utilized to reliably discriminate these two categories (**Vilankar et al. 2014; Breuil et al., 2019**). Therefore, we measured some of these same quantities from our image patches at 40×40 resolution to provide basic descriptive statistics for the shadow and occlusion edges in our databases, and to determine the extent to which such simple statistics could be reliably exploited for image classification purposes.

From both the shadow and occlusion edges, we measured two quantities from linearized RGB versions of the images that were first converted to grayscale: (1) RMS contrast, and (2) Michelson contrast. Conversion to grayscale was accomplished using the standard formula

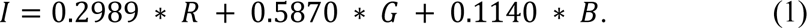

The RMS contrast of the grayscale image *I* is defined as

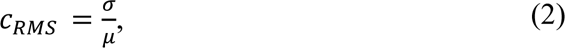

where 𝜎 is the standard deviation of the pixel intensities, and 𝜇 is the mean pixel intensity. Since our selection method only chose image patches which could be cleanly divided into two regions of approximately equal size (see **DiMattina et al. 2012** for details), we were also able to measure the Michelson contrast, which is defined as

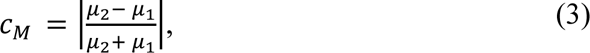

where 𝜇_1_, 𝜇_2_ denote the mean pixel intensity in each of the two regions.

Making use of the MATLAB machine learning toolbox function `fitclinear.m`, we trained a logistic regression classifier (**Bishop, 2006**) on these two variables to see to what extent they could distinguish the two image categories. This was done in two different ways. In the first analysis, we broke each image set (shadows, occlusions) into 5 blocks of non-overlapping images.

The images in **oset-1**, **oset-2** and **shad** comprising each block are listed in **Supplementary Tables S1, S2**. We then trained the classifier on 4 of the blocks (*training sets,* ∼ **50K** images) while testing on the 5^th^ (*test set,* ∼**12K** images). We obtained an estimate of the classifier ability to generalize to novel images by averaging test set performance over all 5 choices of test set. In a second analysis, we trained on N = **16K** images sampled uniformly (with replacement) from all of the images. We then tested classifier performance on a test set comprised of N = **4K** images also sampled uniformly with replacement from all the images.

#### Spatial frequency analysis

In addition to contrast statistics, another potential cue for distinguishing edges caused by shadows from those caused by occlusion boundaries is spatial frequency content. Due to penumbral blur, many (but not all) shadows exhibit a gradual luminance change at their boundary (**Casati & Cavanagh, 2019**), whereas occlusions typically give rise to a relatively sharp luminance transition. Furthermore, psychophysical experiments using classification image techniques have demonstrated that human observers can reliably discriminate sharp luminance edges from blurry ones (**McIlhagga & May, 2012**), suggesting penumbral blur can be potentially employed for shadow identification. We applied the standard FFT to the 40×40 image patches from both categories, which were first pre-processed to have zero mean and identical energy. We then quantified the proportion of stimulus energy in various high spatial-frequency regions of the spectrum. This was accomplished by taking the rotational sum of the power spectrum and then taking the ratio 𝜋_ℎ_ of power above 10 cycles/image to total stimulus power (Nyquist frequency for 40 x 40 images is 20 cycles/image). This parameter 𝜋_ℎ_was combined with the two contrast parameters 𝑐_𝑅𝑀𝑆_, 𝑐_𝑀_to reimplement the logistic regression classifier analysis described in the previous section with a larger feature set.

### Image-computable classifier analyses

Three categories of image-computable machine classifiers were implemented in Python using Keras and Tensorflow (**Chollet, 2021**). To align the images to improve classification accuracy, image patches were rotated into a vertical orientation, and reflected into sine phase (if necessary) so that the left side was always darker than the right side.

As a robust test of model generalization ability, image-computable models were trained on 4 blocks of images (training sets, ∼**50K** images) and tested on a 5^th^ block (test sets, ∼**12K** images) having no overlap with the training sets. This was repeated for a wide range of regularization hyper-parameter values (log_10_ 𝜆 = −3, −2, −1,0,1,2,3), and we report the average performance over 5 test sets for the best hyper-parameter. In our analyses, we found that for all models the performance was generally quite robust over a large range of hyper-parameter values. For exploring model performance characteristics, and comparing model performance with human performance, we trained the models using our full set of images. Therefore, for these analyses we trained models on sets of **16K** (**PIX**, **GFB** models) or **56K** (**FRF** models) images sampled uniformly with replacement from the entire dataset, and tested the models on much smaller sets of **4K** images sampled uniformly from the entire dataset.

#### Gabor filter bank classifier

The second model, which we will refer to as the Gabor Filter Bank (**GFB**) model, is illustrated schematically in **Fig. 4a**. In this model, a fixed set of multiscale Gabor filters (8, 16, 32 pixels) with spatial frequency bandwidths resembling V1 simple cells (**Kovesi, 2000**; **Field, 1987**) was convolved with an input image (40×40 pixels). At each scale, the set of oriented filters includes 6 orientations, 2 phases (even/odd), and 2 contrast polarities (+/-), for a total of 24 oriented filters (**Supplementary Fig. S3)**. The outputs of these filters were half-wave linear rectified (*relu*) and then down-sampled from 40×40 to 5×5 using MAX pooling (**Goodfellow, Bengio & Courville, 2006**). This yields a representation having 1800 dimensions. Binomial logistic regression (**Bishop, 2006**) was performed on these 1800 pooled filter responses as predictor variables with stimulus category (0 = shadow, 1 = occlusion) as the response variable.

#### Filter-rectify-filter classifier

Finally, we implemented the Filter-Rectify-Filter (**FRF**) model often used to account for texture segmentation (**Landy & Graham, 2004; Victor et al., 2017**). This model is illustrated in **Fig. 4b**. As with the **GFB** model, the image was initially convolved with a set of fixed Gabor filters, and the linearly rectified (*relu*) outputs down-sampled using MAX pooling. However, these down-sampled outputs were fed into a three-layer neural network classifier having N = 4, 6, or 10 hidden units with *relu* gain. For analyzing the effects of texture removal on the FRF model (**Fig. 7**), occlusions having vertical or near-vertical orientation were chosen from each image database from a set of images not used for model training. Texture information was removed from these patches using the same method as in our previous work (**DiMattina et al., 2012**), by simply setting the intensity to each pixel in a region equal to the mean intensity over the region. Image patches manipulated in this manner are shown in **Fig. 7a**. This manipulation removes all texture cues while leaving Michelson contrast (**Eq. 2**) unchanged.

### Psychophysical experiments

#### Participants

Participants in this study consisted of N = 18 undergraduate students attending Florida Gulf Coast University enrolled in Psychology courses PSB-4002 (“Brain and Behavior”) and EXP-3202 (“Sensation and Perception”) during the Fall 2021 semester. All participants in these two courses were given the opportunity to complete three experiments (**QT-1**, **QT-2**, **QT-3**) implemented as web surveys in return for extra credit in the course. Before each experiment, participants were required to review and accept an informed consent form. All procedures were approved beforehand by the Institutional Review Board of Florida Gulf Coast University (Protocol 2021-78), in accordance with the Declaration of Helsinki.

#### Procedure

The visual experiments were implemented as a web survey using Qualtrics, an online web survey program (https://www.qualtrics.com/). Participants accessed the link for the Qualtrics survey using their personal computers or smartphones. Image patches used in the Qualtrics surveys used in this study were selected from the McGill Calibrated Color Image Database (**Olmos & Kingdom, 2004**) and then converted to grayscale. In each survey, participants were shown a 40×40 image patch (resized to 96×96, in order to subtend roughly 2-3 degrees of visual angle at normal viewing distances) and asked to indicate whether this patch contained an occlusion edge or a shadow edge. Each survey contained 200 images, half of which belonged to each category, presented in random order. After their response, feedback (correct/incorrect) was provided before presenting the next stimulus. A schematic diagram of the experimental procedure is shown in **Fig. 8a**, and an example survey question is shown in **Supplementary Fig. S11**. For inclusion in the final analysis, students had to complete all three surveys in the correct order, have no blank responses, and perform better than chance (114 or more correct out of 200) on the final 2 surveys, indicating that they had obtained proficiency in the task. Although there were 37 participants who completed at least one survey, only 18 participants met the criteria for inclusion in the final analysis, mostly due to the fact that many participants failed to complete all three surveys.

## Acknowledgements

We would like to thank Curtis Baker for comments on the manuscript, and FGCU graduates Michele DeAngelis and Lauren Anderson for help with developing the database.

