## Supplementary Material for "Distinguishing shadows from surface boundaries using local achromatic cues"

### SUPPLEMENTARY TABLES

| block | edge | shad | Test (2D) | Train (2D) | Test (3D) | Train (3D) |
|---|---|---|---|---|---|---|
| 1 | 1-20 | 1-9 | 0.629 | 0.702 | 0.534 | 0.711 |
| 2 | 21-40 | 10-18 | 0.567 | 0.726 | 0.585 | 0.725 |
| 3 | 41-60 | 19-27 | 0.599 | 0.708 | 0.603 | 0.701 |
| 4 | 61-80 | 28-36 | 0.800 | 0.677 | 0.808 | 0.669 |
| 5 | 81-100 | 37-45 | 0.790 | 0.681 | 0.790 | 0.681 |
| ***AVG*** | - | - | ***0.677*** | ***0.699*** | ***0.664*** | ***0.697*** |

**Supplementary Table S1:** Logistic regression classifier analysis for **oset-1**. Each block used the indicated images as the test set and trained on the remainder. This ensured no overlap between the test and training sets. The 2D in parentheses refers to classification using only Michelson and RMS Contrast parameters. The 3D analyses also include a parameter measuring proportion of stimulus energy in high spatial frequencies.

| block | edge | shad | Test (2D) | Train (2D) | Test (3D) | Train (3D) |
|---|---|---|---|---|---|---|
| 1 | 1-4 | 1-9 | 0.809 | 0.677 | 0.797 | 0.667 |
| 2 | 5-8 | 10-18 | 0.664 | 0.713 | 0.644 | 0.711 |
| 3 | 9-12 | 19-27 | 0.609 | 0.722 | 0.603 | 0.724 |
| 4 | 13-16 | 28-36 | 0.740 | 0.691 | 0.663 | 0.697 |
| 5 | 17-20 | 37-45 | 0.601 | 0.730 | 0.590 | 0.728 |
| ***AVG*** | - | - | ***0.685*** | ***0.707*** | ***0.659*** | ***0.705*** |

**Supplementary Table S2:** Same as **Supplementary Table S1** but for occlusions taken from **oset-2**.

| block | edge | shad | Test | Train |
|---|---|---|---|---|
| 1 | 1-20 | 1-9 | 0.8448 | 0.8318 |
| 2 | 21-40 | 10-18 | 0.8342 | 0.8498 |
| 3 | 41-60 | 19-27 | 0.7483 | 0.8531 |
| 4 | 61-80 | 28-36 | 0.7268 | 0.8615 |
| 5 | 81-100 | 37-45 | 0.7715 | 0.8410 |
| ***AVG*** | - | - | ***0.7851*** | ***0.8474*** |

**Supplementary Table S3:** Gabor Filter Bank (**GFB**) classifier analysis for **oset-1** at the best hyper-parameter value. Each block used the indicated images as the test set and trained on the remainder. Performance was robust across a wide range of hyper-parameters (**Supplementary Fig. S12**)

| block | edge | shad | Test | Train |
|---|---|---|---|---|
| 1 | 1-4 | 1-9 | 0.6374 | 0.7972 |
| 2 | 5-8 | 10-18 | 0.6713 | 0.8097 |
| 3 | 9-12 | 19-27 | 0.7519 | 0.8513 |
| 4 | 13-16 | 28-36 | 0.6884 | 0.8629 |
| 5 | 17-20 | 37-45 | 0.7922 | 0.8180 |
| ***AVG*** | - | - | ***0.7082*** | ***0.8278*** |

**Supplementary Table S4:** Same as **Supplementary Table S3**, but for **oset-2**.

| Set | GFB | | FRF-4 | | FRF-6 | | FRF-10 | |
|---|---|---|---|---|---|---|---|---|
| | ***os-1*** | ***os-2*** | ***os-1*** | ***os-2*** | ***os-1*** | ***os-2*** | ***os-1*** | ***os-2*** |
| **1** | 0.798 | 0.791 | 0.875 | 0.886 | 0.884 | 0.889 | 0.883 | 0.898 |
| **2** | 0.800 | 0.820 | 0.879 | 0.923 | 0.888 | 0.910 | 0.902 | 0.930 |
| **3** | 0.810 | 0.800 | 0.874 | 0.895 | 0.889 | 0.894 | 0.902 | 0.904 |
| **4** | 0.789 | 0.804 | 0.865 | 0.878 | 0.879 | 0.888 | 0.888 | 0.892 |
| **AVG** | ***0.799*** | ***0.804*** | ***0.873*** | ***0.896*** | ***0.885*** | ***0.895*** | ***0.894*** | ***0.906*** |

**Supplementary Table S5:** Classifier performance on 4 test sets not used for training models. In this analysis, both the training sets and the test sets were sampled uniformly with replacement from all the images. Optimized hyper-parameters were used to train each model (**Supplementary Fig. S5**).

| block | edge | shad | Test | Train |
|---|---|---|---|---|
| 1 | 1-20 | 1-9 | 0.8834 | 0.8424 |
| 2 | 21-40 | 10-18 | 0.8420 | 0.8606 |
| 3 | 41-60 | 19-27 | 0.7568 | 0.8742 |
| 4 | 61-80 | 28-36 | 0.7317 | 0.8780 |
| 5 | 81-100 | 37-45 | 0.8021 | 0.8589 |
| ***AVG*** | - | - | ***0.8031*** | ***0.8628*** |

**Supplementary Table S6:** Filter-Rectify-Filter model with 6 hidden units (**FRF-6**) classifier analysis for **oset-1** at the best hyper-parameter value. Each block used the indicated images as the test set and trained on the remainder. Performance was robust across a wide range of hyper-parameters.

| block | edge | shad | Test | Train |
|---|---|---|---|---|
| 1 | 1-4 | 1-9 | 0.7065 | 0.8909 |
| 2 | 5-8 | 10-18 | 0.7240 | 0.8795 |
| 3 | 9-12 | 19-27 | 0.7381 | 0.9002 |
| 4 | 13-16 | 28-36 | 0.7171 | 0.8955 |
| 5 | 17-20 | 37-45 | 0.8020 | 0.8844 |
| ***AVG*** | - | - | ***0.7375*** | ***0.8901*** |

**Supplementary Table S7:** Same as **Supplementary Table S6**, but for **oset-2**.

| block | edge | shad | Test | Train |
|---|---|---|---|---|
| 1 | 1-20 | 1-9 | 0.8807 | 0.8377 |
| 2 | 21-40 | 10-18 | 0.8379 | 0.8501 |
| 3 | 41-60 | 19-27 | 0.7676 | 0.8613 |
| 4 | 61-80 | 28-36 | 0.7169 | 0.8744 |
| 5 | 81-100 | 37-45 | 0.7934 | 0.8374 |
| ***AVG*** | - | - | ***0.7993*** | ***0.8522*** |

**Supplementary Table S8:** Filter-Rectify-Filter model with 4 hidden units (**FRF-4**) classifier analysis for **oset-1** at the best hyper-parameter value.

| block | edge | shad | Test | Train |
|---|---|---|---|---|
| 1 | 1-4 | 1-9 | 0.6826 | 0.8234 |
| 2 | 5-8 | 10-18 | 0.7052 | 0.7923 |
| 3 | 9-12 | 19-27 | 0.7535 | 0.7989 |
| 4 | 13-16 | 28-36 | 0.7069 | 0.8287 |
| 5 | 17-20 | 37-45 | 0.8246 | 0.7811 |
| ***AVG*** | - | - | ***0.7346*** | ***0.8049*** |

**Supplementary Table S9:** Same as **Supplementary Table S8**, but for **oset-2**.

| block | edge | shad | Test | Train |
|---|---|---|---|---|
| 1 | 1-20 | 1-9 | 0.8641 | 0.8577 |
| 2 | 21-40 | 10-18 | 0.8488 | 0.8702 |
| 3 | 41-60 | 19-27 | 0.7588 | 0.8786 |
| 4 | 61-80 | 28-36 | 0.7406 | 0.8911 |
| 5 | 81-100 | 37-45 | 0.8109 | 0.8726 |
| ***AVG*** | - | - | ***0.8046*** | ***0.8741*** |

**Supplementary Table S10:** Filter-Rectify-Filter model with 4 hidden units (**FRF-10**) classifier analysis for **oset-1** at the best hyper-parameter value.

| block | edge | shad | Test | Train |
|---|---|---|---|---|
| 1 | 1-4 | 1-9 | 0.7053 | 0.9096 |
| 2 | 5-8 | 10-18 | 0.7352 | 0.9081 |
| 3 | 9-12 | 19-27 | 0.7231 | 0.9240 |
| 4 | 13-16 | 28-36 | 0.7209 | 0.9078 |
| 5 | 17-20 | 37-45 | 0.7966 | 0.9113 |
| ***AVG*** | - | - | ***0.7362*** | ***0.9122*** |

**Supplementary Table S11:** Same as **Supplementary Table S10**, but for **oset-2**.

| | **occlusion** | | **shadow** | |
|---|---|---|---|---|
| | ***oset-1*** | ***oset-2*** | ***oset-1*** | ***oset-2*** |
| **FRF-4** | 0.57 | 0.93 | -2.51 | -1.92 |
| **FRF-6** | 0.72 | 2.07 | -2.33 | -2.41 |
| **FRF-10** | 0.36 | 1.21 | -2.68 | -2.34 |

**Supplementary Table S12**.: Median values of $\Delta$ for occlusions classified as occlusions or incorrectly classified as for all **FRF** models (**oset-1**: N = 191; **oset-2**: N = 596) Median values were significantly different from zero with $p < 0.001$ (Wilcoxon rank-sum test).

#### SUPPLEMENTARY FIGURE CAPTIONS

**Supplementary Figure S1:** Same as **Fig. 2a** in main text but for a subset of 19 images in **oset-1** which were analyzed by **Vilankar et al. (2014).**

**Supplementary Figure S2**: Same as **Fig. 2a** in main text but for image patches which were normalized to the range [0, 1].

**Supplementary Figure S3**: Gabor filters at one spatial scale (16x16) used in both the **GFB** model and **FRF** models (**Fig. 6**).

**Supplementary Figure S4: GFB** model performance at generalizing to novel test sets comprised of 20% of the images (not overlapping with 80% used for training) for various values of the regularization hyperparameter lambda. Top panels show analysis for **oset-1**, bottom for **oset-2**. Left panels show test set performance, right panels show training set performance. Dots indicate individual folds, the blue curve denotes average performance.

**Supplementary Figure S5**: Results of cross validation analysis for all 3 models (**GFB**, **FRF**) when both trained and tested on sets sampled uniformly from all of the images. Average prediction performance on the test folds is plotted (blue curve) as a function of regularization hyper-parameter value (lambda). Blue dots show results for individual folds.

**Supplementary Figure S6:** Stimuli that maximize + minimize the occlusion probability predicted by the **GFB** and **FRF** models trained to distinguish shadows from occlusions (**oset-1**, **oset-2**).

(**a**) *Left:* Stimuli obtained via numerical optimization with highest predicted occlusion probability by the **GFB** model. We see consistent results for both training sets, and over multiple optimization runs. *Right:* Same as left column, but with highest predicted shadow probability by the **GFB** model (equivalently, lowest predicted occlusion probability).

(**b**) Same as (a) but for **FRF-4**.

(**c**) Same as (a) but for **FRF-6**.

(**d**) Same as (a) but for **FRF-10**.

**Supplementary Figure S7**: Same as **Supplementary Fig. S4** but for **FRF-6**.

**Supplementary Figure S8**: Same as **Supplementary Fig. S4** but for **FRF-4**.

**Supplementary Figure S9**: Same as **Supplementary Fig. S4** but for **FRF-10**.

**Supplementary Figure S10**: Optimal stimuli for the hidden units in the **FRF** models with 4 and 6 hidden units trained with both sets of occlusions (**oset-1**, **oset-2**).

(**a**) **FRF-4** hidden unit receptive fields. Sign of output weight is indicated with a (+) or (-).

(**b**) Same as (a) but for **FRF-6**.

**Supplementary Figure S11**: Example question from the Qualtrics surveys.

**Supplementary Figure S12**: Distribution of SDT decision threshold $\gamma$ for each individual survey.

**Supplementary Figure S13:** Probability of classification as an occlusion by human observers (**AO**) and each model (**GFB**, **FRF**) for all N = 200 edges in each survey. Green symbols indicate occlusions, magenta indicates shadows.

**Supplementary Fig. S1**

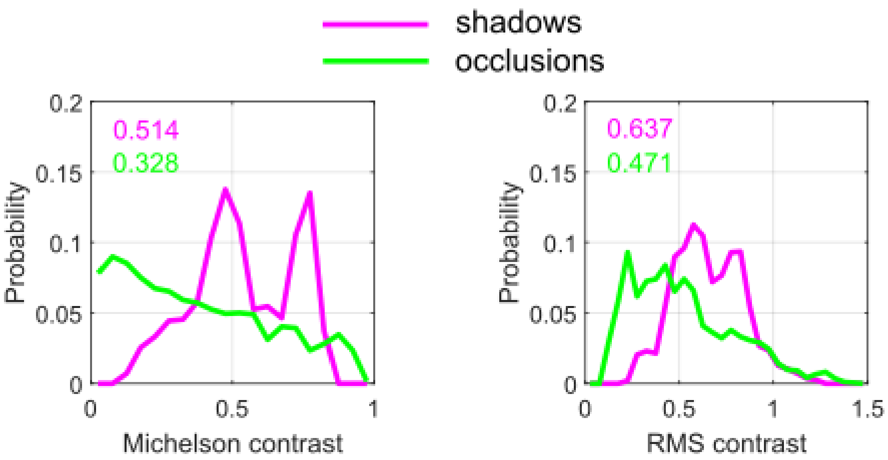

**Supplementary Fig. S2**

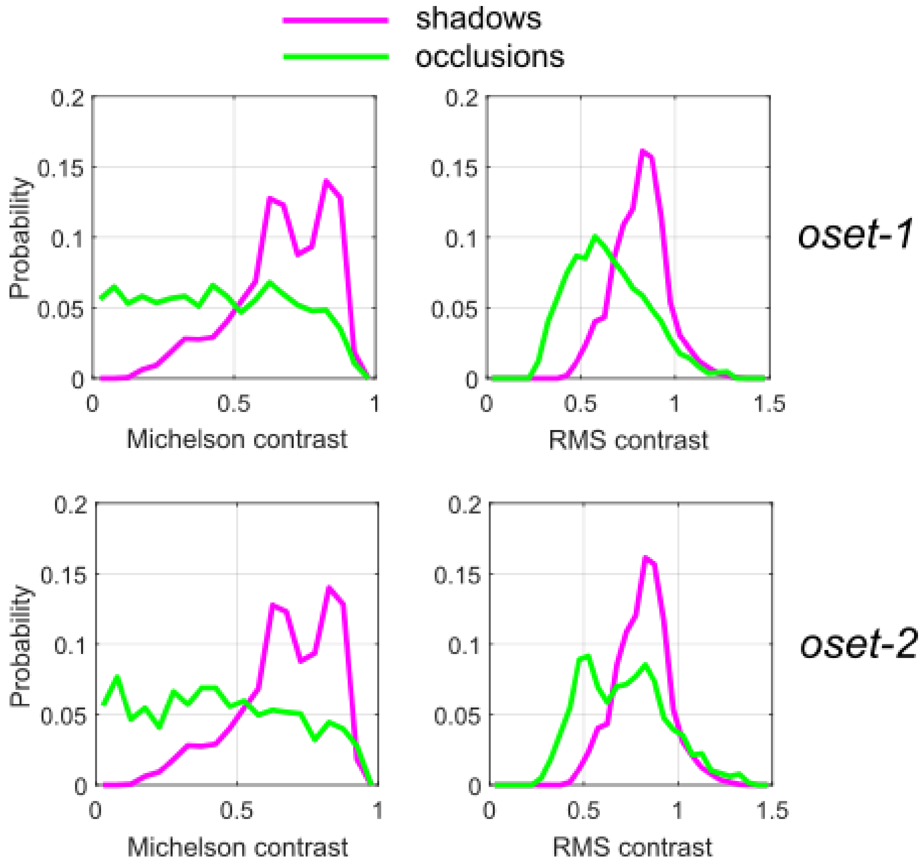

**Supplementary Fig. S3**

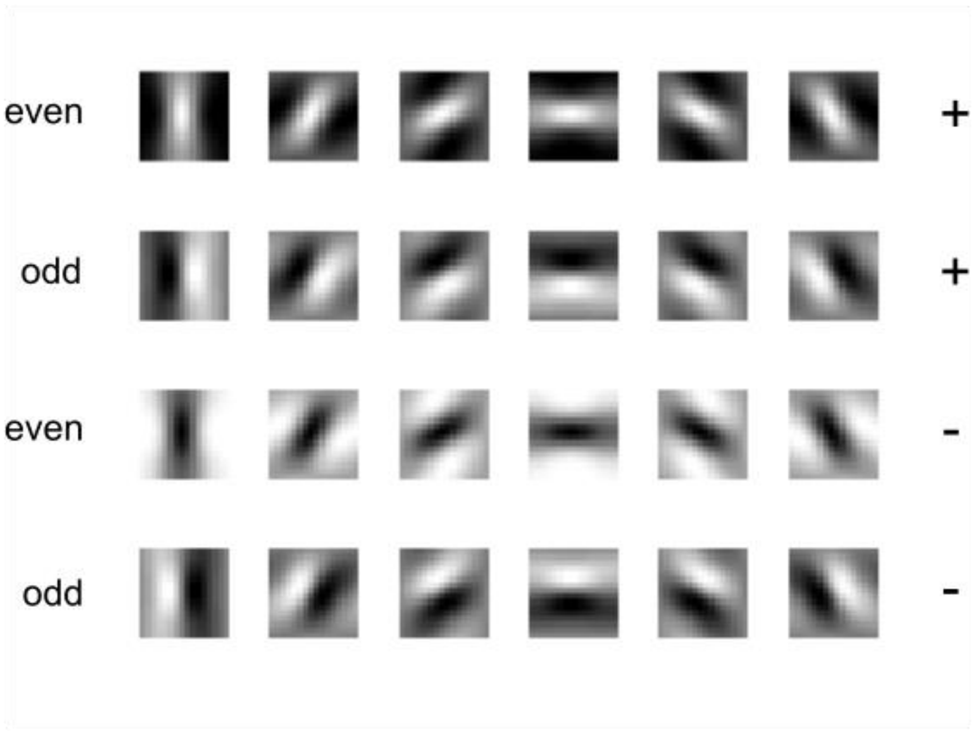

**Supplementary Fig. S4**

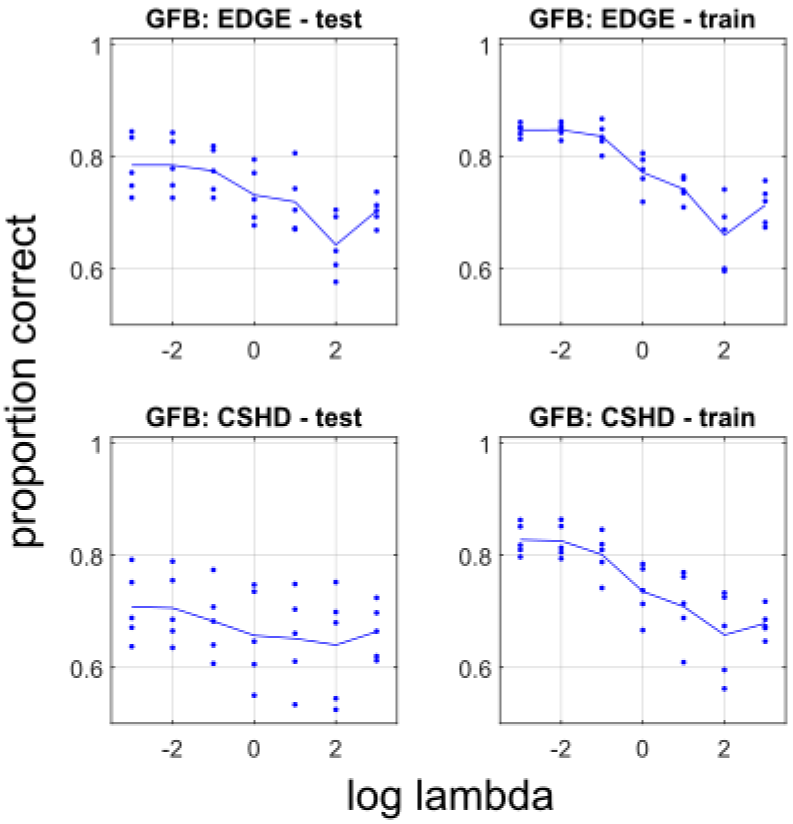

**Supplementary Fig. S5**

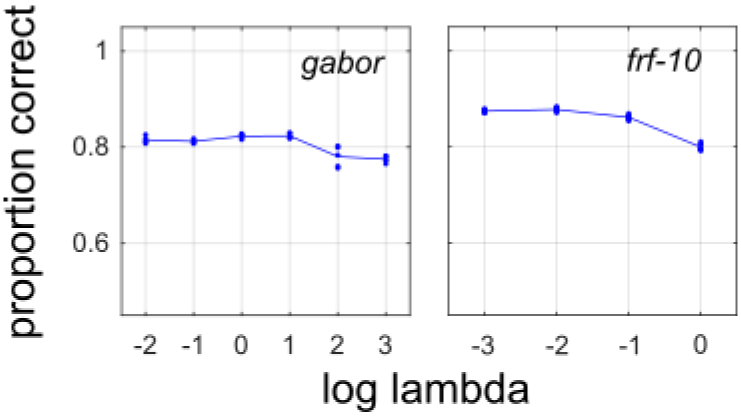

**Supplementary Fig. S6**

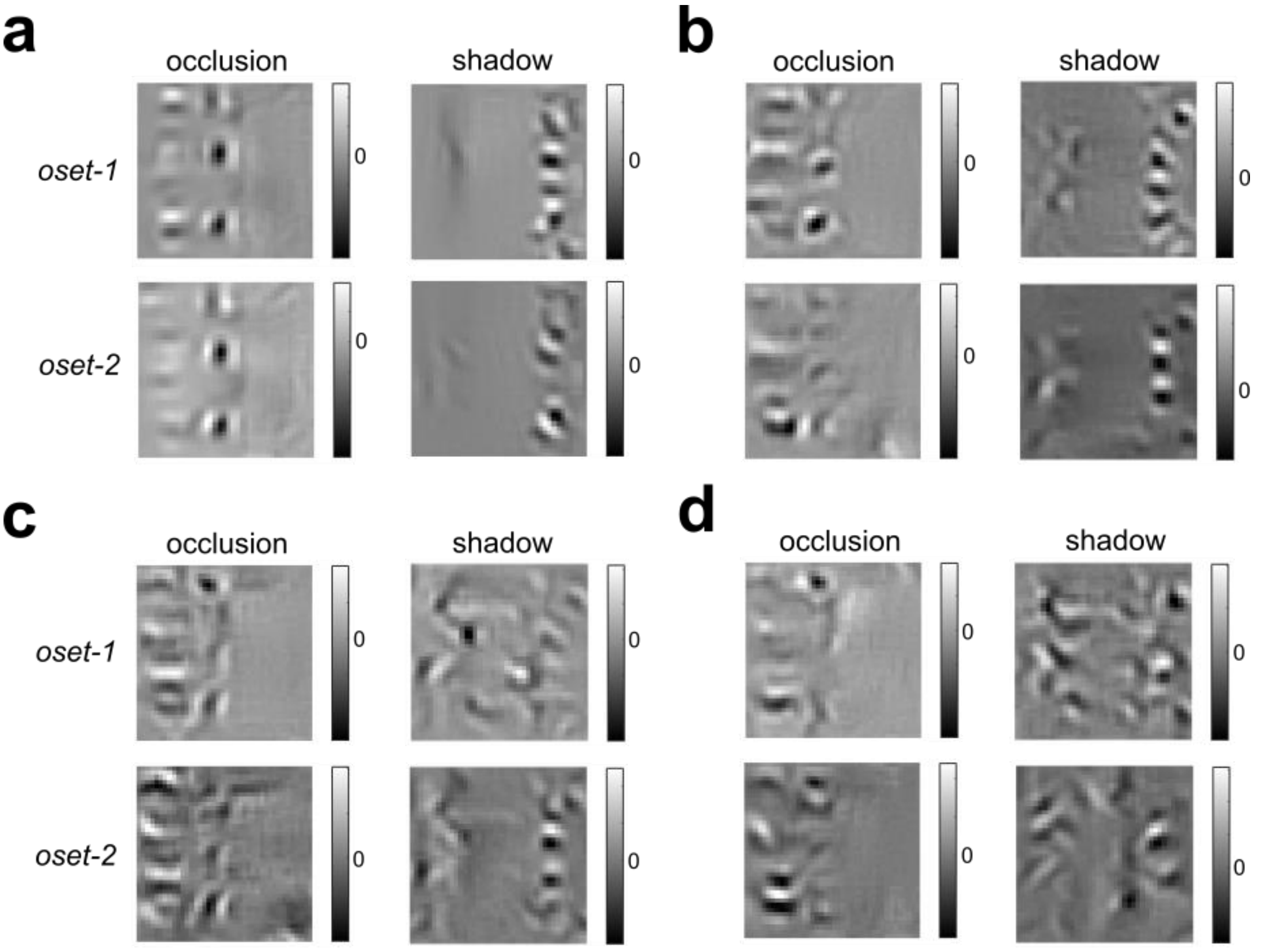

**Supplementary Fig. S7**

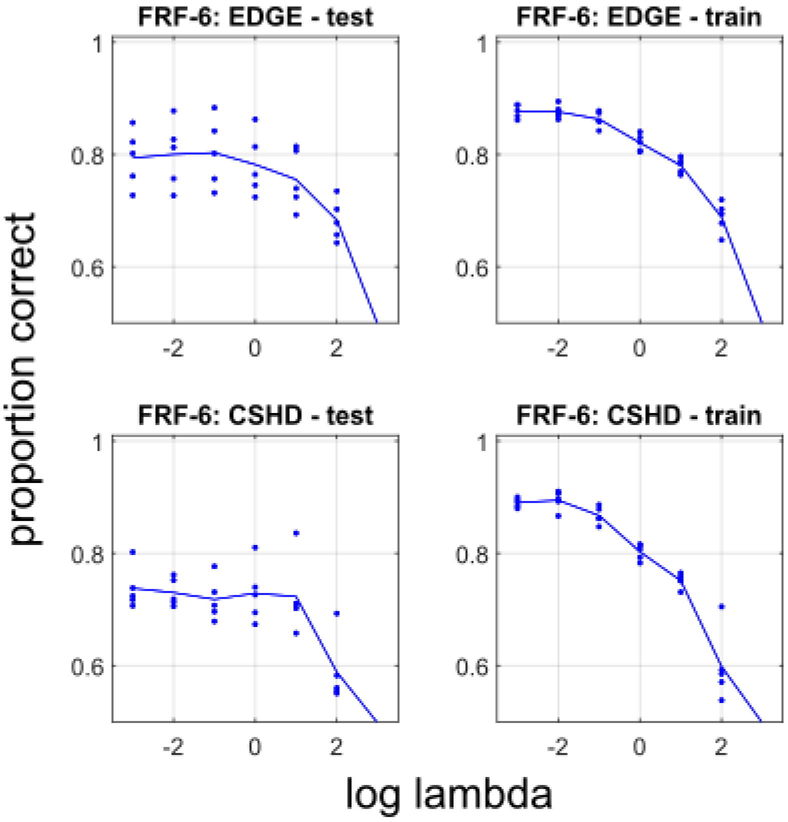

**Supplementary Fig. S8**

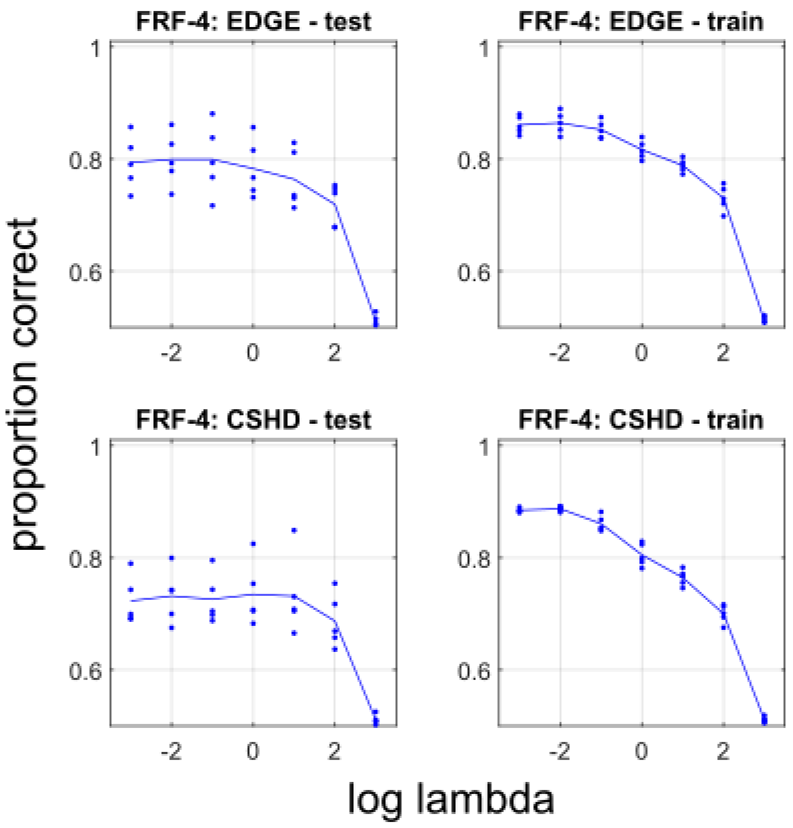

**Supplementary Fig. S9**

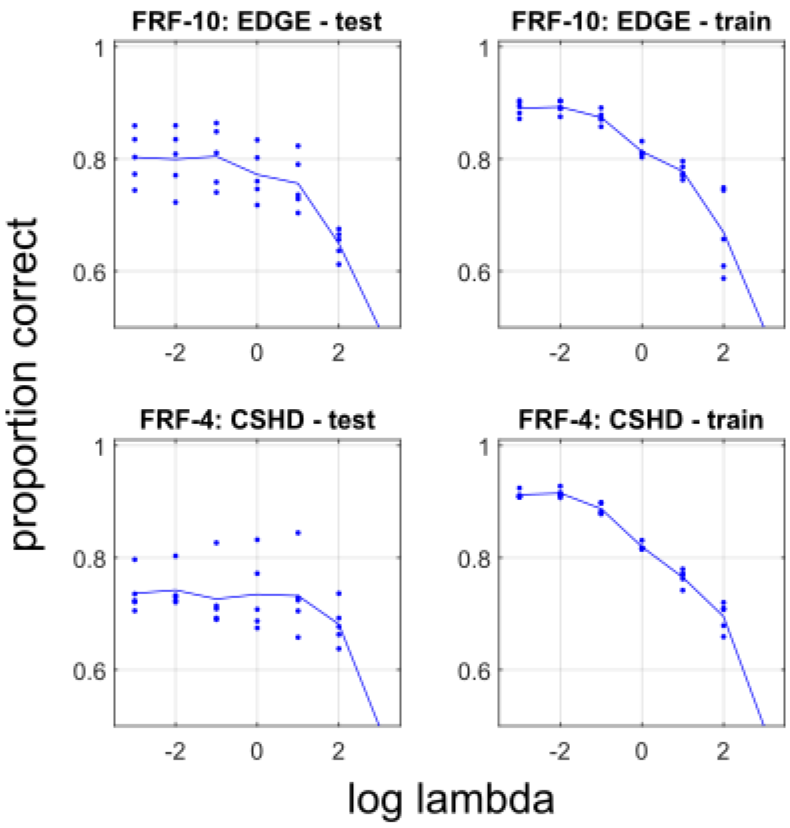

**Supplementary Fig. S10**

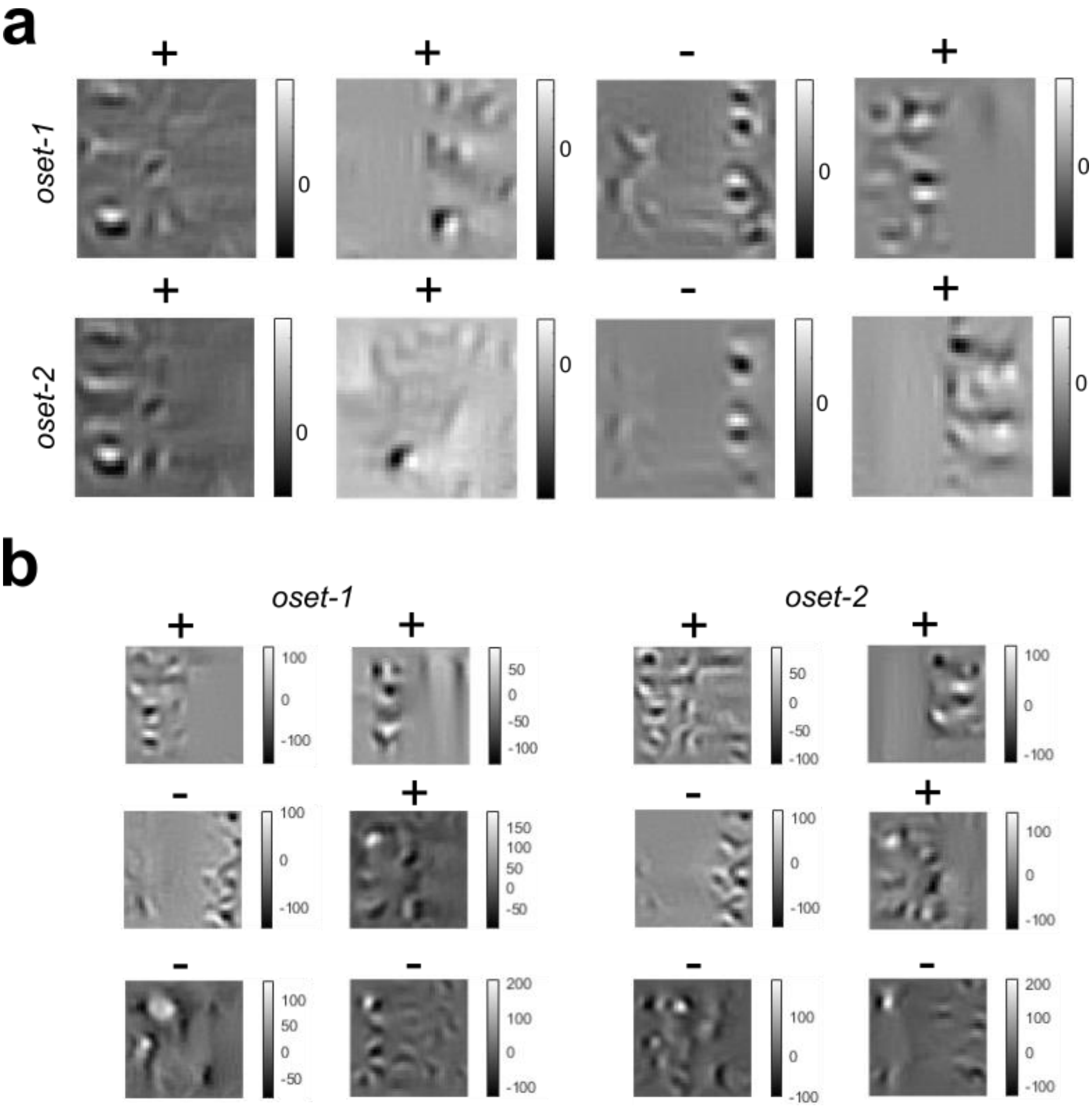

**Supplementary Fig. S11**

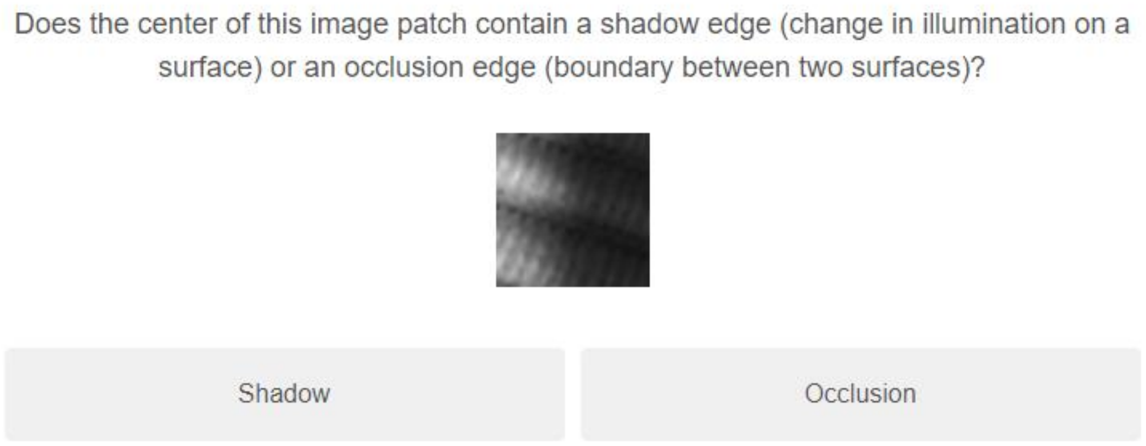

**Supplementary Fig. S12**

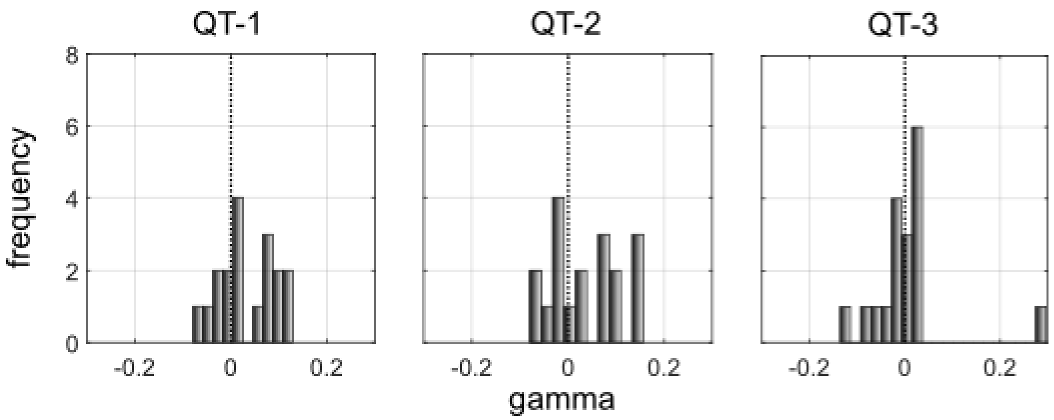

**Supplementary Fig. S13**

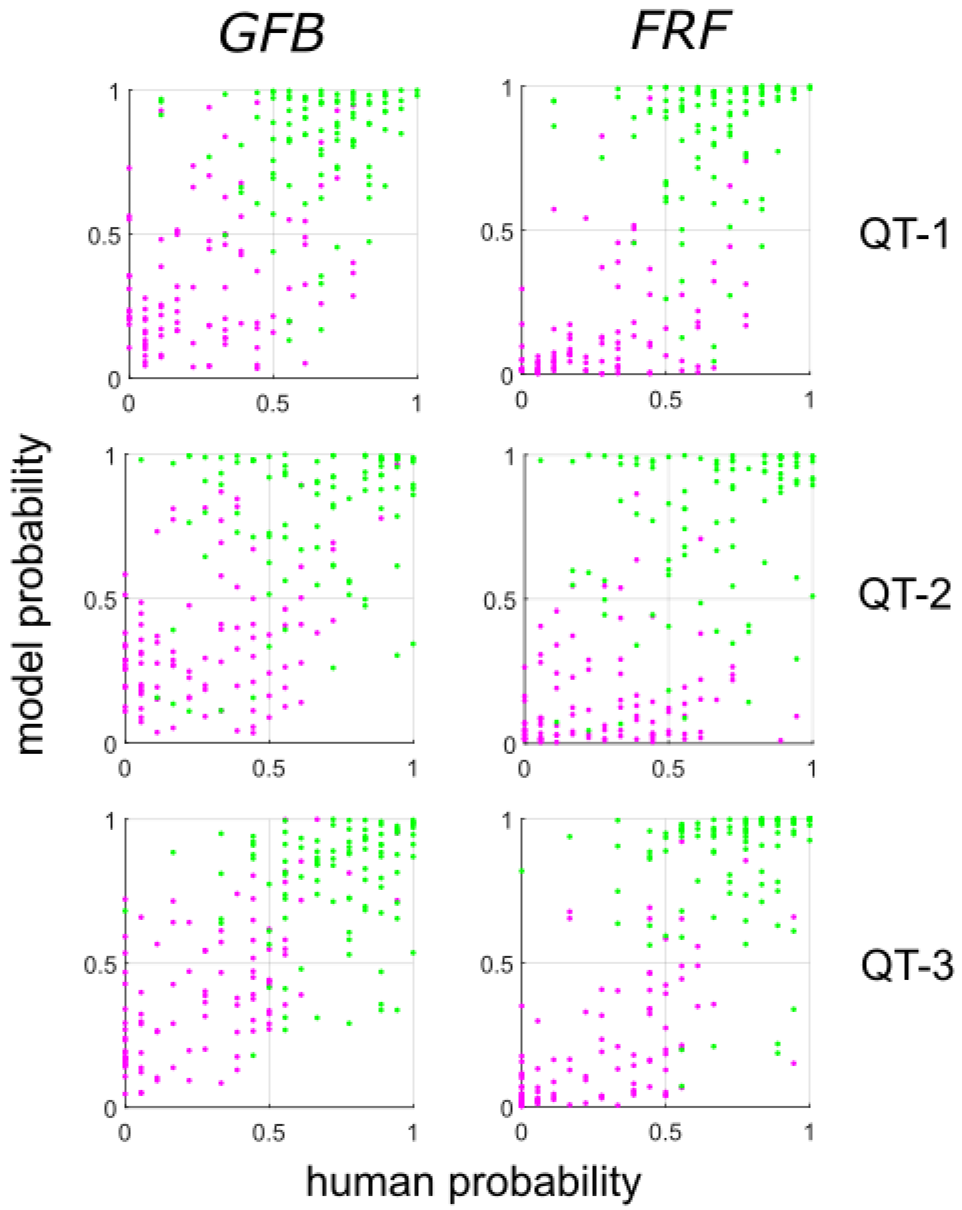
